# Lymph node expansion predicts magnitude of vaccine immune response

**DOI:** 10.1101/2022.10.25.513749

**Authors:** Alexander J. Najibi, Ryan S. Lane, Miguel C. Sobral, Benjamin R. Freedman, Joel Gutierrez Estupinan, Alberto Elosegui-Artola, Christina M. Tringides, Maxence O. Dellacherie, Katherine Williams, Sören Müller, Shannon J. Turley, David J. Mooney

## Abstract

Lymph nodes (LNs) dynamically expand in response to immunization, but the relationship between LN expansion and the accompanying adaptive immune response is unclear. Here, we first characterized the LN response across time and length scales to vaccines of distinct strengths. High-frequency ultrasound revealed that a bolus ‘weak’ vaccine induced a short-lived, 2-fold volume expansion, while a biomaterial-based ‘strong’ vaccine elicited an ∼7-fold LN expansion, which was maintained several weeks after vaccination. This latter expansion was associated with altered matrix and mechanical properties of the LN microarchitecture. Strong vaccination resulted in massive immune and stromal cell engagement, dependent on antigen presence in the vaccine, and conventional dendritic cells and inflammatory monocytes upregulated genes involved in antigen presentation and LN enlargement. The degree of LN expansion following therapeutic cancer vaccination strongly correlated with vaccine efficacy, even 100 days post-vaccination, and direct manipulation of LN expansion demonstrated a causative role in immunization outcomes.

## Introduction

Lymph nodes (LNs) dynamically expand and contract as they orchestrate adaptive immune responses to disease and immunization [1–5]. This process is driven by changes in cellular contractility, proliferation, and lymphocyte retention [2, 6–10]. At rest, fibroblastic reticular cells (FRCs) in the LN stroma remain in a tensile state, providing resistance to LN expansion. However, engagement of dendritic cell (DC)-expressed C-type lectin-like receptor 2 (CLEC-2) with FRC-expressed podoplanin during an immune response reduces stromal tension, enabling cellular elongation and LN expansion [2, 9]. While this impact of cellular contractility in LN expansion has been widely explored, changes in LN tissue-scale mechanical properties following vaccination remain incompletely understood, and may inform cellular behavior [11, 12]. Further, a successful immune response engages a variety of immune and stromal cell types, culminating in T and B cell proliferation, germinal center formation, and antibody production. However, the time-dependent cellular and transcriptional profiles within LNs leading to differential outcomes based on vaccine formulation remain unclear. With a diversity of novel vaccine strategies under exploration [13], discerning these cellular mechanisms could support vaccine development.

In this work, we assessed the cellular and tissue-scale LN responses to ‘strong’ and ‘weak’ vaccines with distinct therapeutic outcomes. Vaccine-draining LNs (dLNs) were imaged longitudinally using high frequency ultrasound (HFUS), a simple, noninvasive technique for visualizing superficial internal structures with high (∼20μm) axial resolution, to establish the magnitude and kinetics of LN response [14]. Gross LN expansion was then analyzed in comparison to outcomes of immunization (i.e. antigen-specific T cell and antibody responses, therapeutic antitumor efficacy), tissue-scale mechanical and matrix properties, cellular composition, and transcriptional signatures. Finally, the ability of LN expansion to directly enhance the efficacy of vaccination was explored.

## Results

### Vaccine formulation alters the magnitude and duration of lymph node expansion

First, LN expansion in response to a traditional bolus (liquid) vaccine was compared with a mesoporous silica (MPS) rod-based vaccine formulation found to elicit robust immune responses against diverse antigen targets [15–18]; these served as models of ‘weak’ and ‘strong’ vaccination, respectively. Draining (dLN; ipsilateral to vaccine site) and non-draining (ndLN; contralateral) inguinal LNs of mice immunized with MPS or bolus vaccines protein were imaged for 100 days post-vaccination using HFUS.

While PBS injection did not affect LN volume, both vaccine formulations induced LN expansion, but to different extents (Fig. 1a-b, Supplementary Fig. 1a-b). The bolus vaccine led to a two-fold, transient increase in LN volume, while the MPS vaccine induced a more substantial (∼7x) LN expansion over one week which was maintained for approximately three weeks. Although LN volume in the MPS-vaccinated mice began to decrease ∼20 days after immunization, it remained elevated out to 100 days (Supplementary Fig. 1c). NdLNs did not change in volume with either vaccine (Fig. 1c).

**Figure 1.**
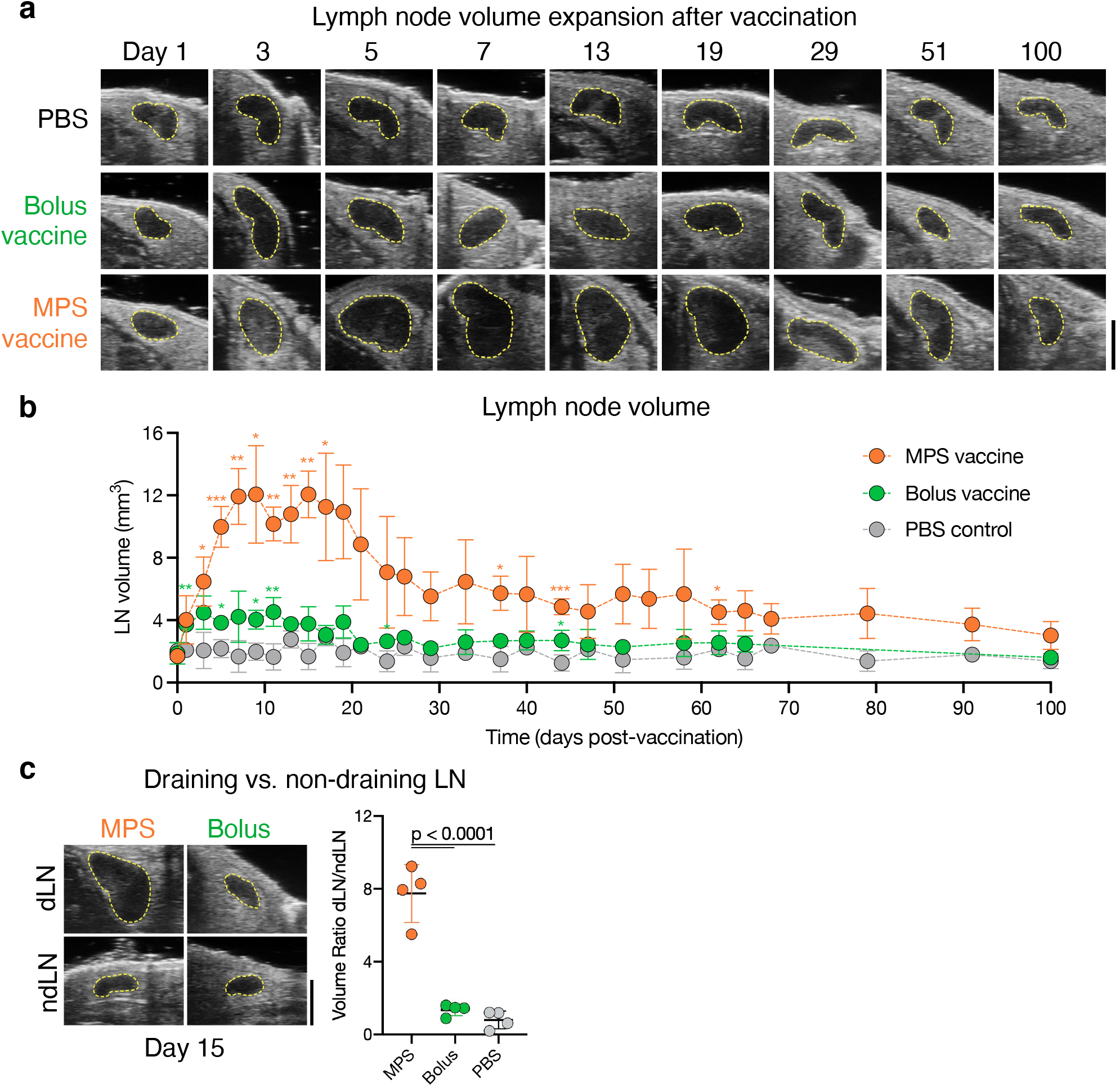
MPS vaccination induces robust, prolonged LN expansion. Mice were immunized with MPS or bolus vaccines delivering GM-CSF, CpG, and OVA protein, and compared to PBS-injected controls. Vaccine-draining and non-draining LNs were longitudinally imaged using high frequency ultrasound (HFUS). (a) Representative HFUS images of vaccine draining LNs out to 100 days after vaccination. (b) Quantification of vaccine draining LN volume over time. Statistical analysis was performed using a two-way ANOVA with repeated measures. Significance relative to the PBS group is depicted at each timepoint (*P < 0.05, ** P < 0.01, ***P < 0.001, ****P < 0.001). (c) Representative HFUS images of MPS or bolus vaccine-draining or non-draining LNs 15 days after vaccination (left) and quantification of dLN/ndLN volume ratio (right). Statistical analysis was performed using analysis of variance (ANOVA) with Tukey’s post hoc test. For a-c, n = 7-8 biologically independent animals per group, imaged longitudinally in two cohorts; means depicted; error bars, s.d.

### LN expansion involves location-specific alterations in lymph node matrix and mechanical properties

LN tissue-scale mechanical properties and extracellular matrix (ECM) distribution were next characterized during expansion, using the MPS vaccine to elicit maximal enlargement. LN collagen architecture was largely maintained following expansion, as expected (Fig. 2a) [6]. Hyaluronic acid (HA) localization was increased in the periphery/follicle, most visibly 7 days after immunization (Fig. 2a-b, Supplementary Fig. 2a). In contrast, the cellular F-actin signal was greater towards the center of both control and MPS dLNs, with greater polarization between the center-periphery in the MPS condition (Fig. 2c, Supplementary Fig. 2b-d). Through nanoindentation of thick (∼500μm) LN slices (Fig. 2d), we found that MPS-vaccinated mouse LNs (day 7) had reduced stiffness (G’) and loss modulus (G’’) compared to control LNs (Fig. 2e-f). Viscoelasticity, measured by G’’/G’ (tan(δ)), was significantly increased in MPS dLNs compared to LNs from control mice, suggesting decreased matrix crosslinking (Fig. 2g).

**Figure 2.**
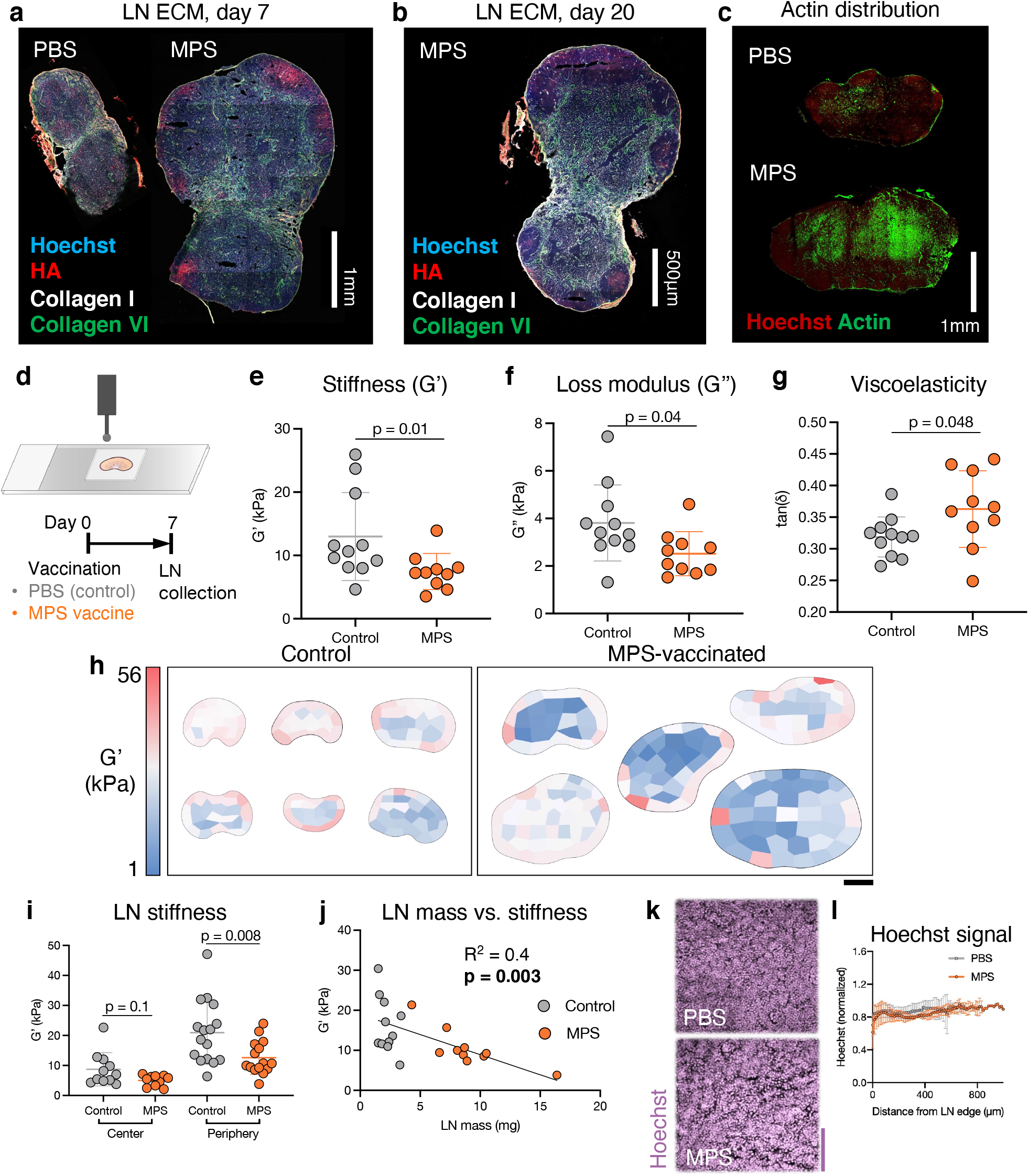
Location-specific alterations in lymph node matrix and mechanics accompany expansion. Mice were treated with MPS vaccines (delivering GM-CSF, CpG, OVA) or PBS, and LNs were harvested after 7 and 20 days. (a) Representative IHC images depicting LN ECM on day 7. (b) Representative IHC image depicting LN ECM on day 20. (c) Representative IHC images of LNs stained for F-actin on day 7. (d) Schematic depicting nanoindentation of a thick LN slice (above) and experimental timeline (below). Mean (e) G’, (f) G’’, and (g) tan(δ) across LNs. Statistical analysis was performed using a Mann-Whitney test (e) or two-tailed t test (f-g). n = 10-11 biologically independent animals per group; results are combined from two independent experiments; means depicted; error bars, s.d. (h) Heatmaps depicting G’ across individual LNs. Scale bar = 1mm. (i) Mean G’ of sample points across each LN, separated into those collected on the center or periphery. n = 10-16 biologically independent animals per group; results are combined from three independent experiments; means depicted; error bars, s.d. Statistical analysis was performed using a Mann-Whitney test. (j) Plot of LN mass versus mean G’. For e, f, g, i, and j, each data point represents a unique LN/mouse. (k) Representative IHC images depicting Hoechst stain within LNs on day 7. Scale bar = 100μm. (l) Quantification of Hoechst signal across LNs; n = 3 biologically independent animals per group.

Spatial variations in mechanics across LNs were next investigated using nanoindentation (Fig. 2h, Supplementary Fig. 3). Both control and MPS-vaccinated mouse LNs were softer and more viscoelastic in the center than the periphery, and this finding was confirmed through intentional sampling on the center or periphery of naïve LNs (Supplementary Fig. 4a-e). The LN periphery (∼12 kPa) was approximately twice as stiff as the center (∼6 kPa). Interestingly, after vaccination, LN G’ and G’’ were only significantly altered on the periphery, while tan(δ) increased only in the LN center (Fig. 2i, Supplementary Fig. 5a-e). LN peripheral stiffness correlated negatively with LN mass, suggesting that the degree of tissue softening relates to the extent of LN enlargement induced by vaccination (Fig. 2j). LN cellular distribution and tissue density remained unaltered, despite expansion (Fig. 2k-l, Supplementary Fig. 6a-d).

### Cellular activity in lymph nodes is regulated by vaccine formulation

Considering tissue-level changes may impact or reflect cellular responses, changes in LN cellularity during expansion were next characterized (Supplementary Fig. 7a, Supplementary Fig. 8). Cellular expansion was greater and more sustained in MPS-vaccinated mice than in the bolus vaccine or PBS-injected control; notably, the total cell count within a LN correlated with its volume (Fig. 3a-b). As early as day 4, monocytes, neutrophils, and macrophages were expanded in MPS dLNs, while conventional DCs, plasmacytoid DCs, and T cells peaked at day 7 before declining over time (Fig. 3c-d, Supplementary Fig. 7b-j). Monocytes in particular expanded ∼80-fold in MPS dLNs compared to PBS controls as early as 4 days after vaccination, compared to ∼25x expansion in the bolus group, and this increase was maintained for several weeks only in the MPS condition (Supplementary Fig. 7p). B cells also significantly expanded by day 7 and remained elevated until day 17 (Fig. 3e). A variety of stromal cells expanded following MPS vaccination, typically peaking later (days 11-17) than the immune cells, with the exception of NK cells, which also tended to expand later (days 7-11) (Fig. 3f, Supplementary Fig. 7k-o). By comparison, changes in the bolus vaccine group were more modest, and PBS vaccine dLNs and ndLNs from all groups demonstrated minimal changes in cell populations.

**Figure 3.**
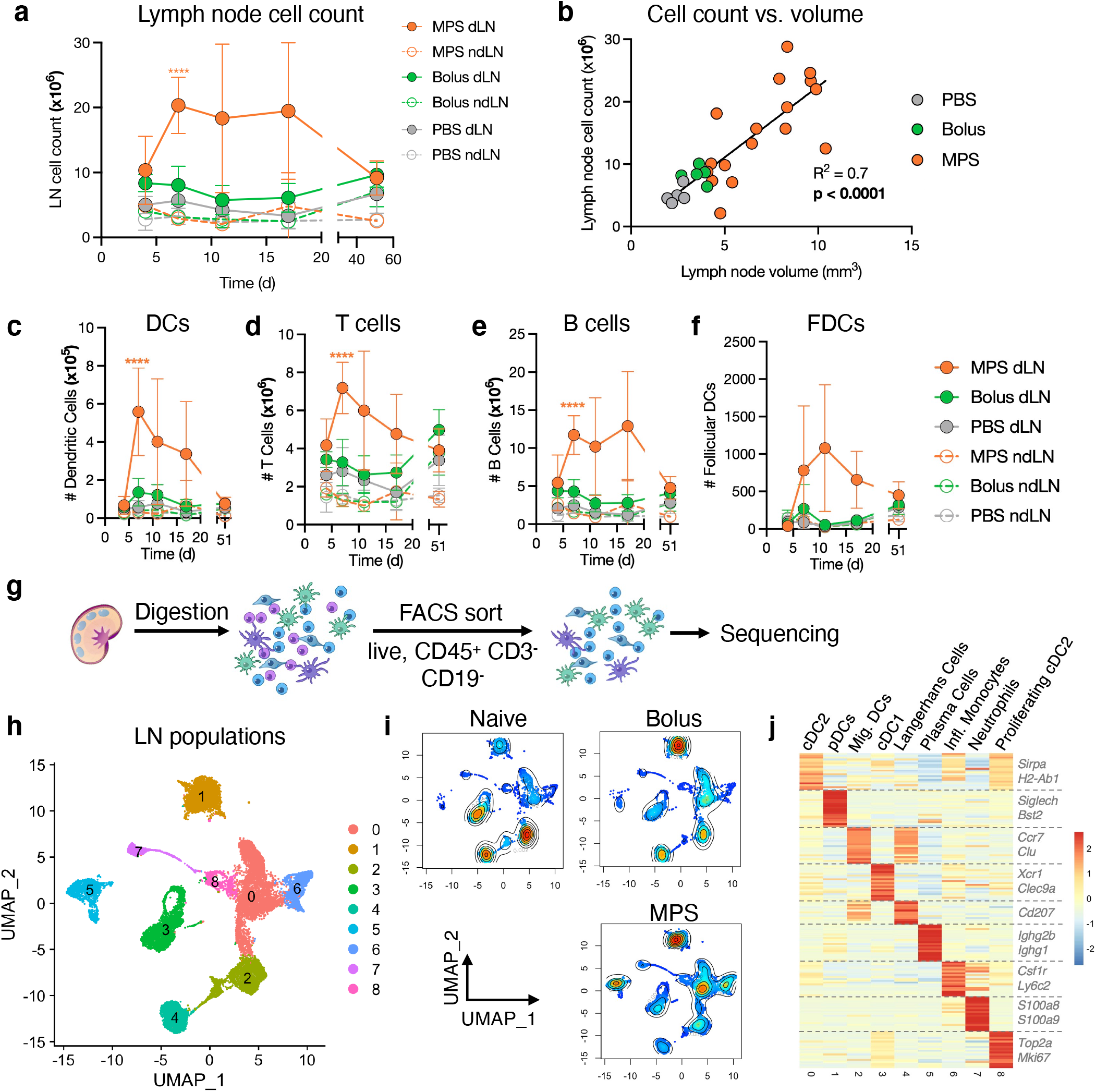
MPS vaccination alters cellularity of lymph nodes. Mice were immunized with MPS or bolus vaccines containing GM-CSF, CpG, and OVA protein, euthanized on days 4, 7, 11, 17, and 51 for LN harvest and analysis through flow cytometry, and compared to PBS-injected controls. (a) Total LN cell counts over time. (b) Linear regression of LN cell count on a given day versus volume (measured through HFUS). Numbers of (c) dendritic cells, (d) T cells, (e) B cells, and (f) follicular dendritic cells over time. For a and c-f, n = 4-5 biologically independent animals per group per timepoint; means depicted; error bars, s.d. Statistical analysis was performed using analysis of variance (ANOVA) with Tukey’s post hoc test for normally distributed samples, and a Kruskal-Wallis test with Dunn’s post hoc test otherwise; only differences present between one group and all other groups are shown (*P < 0.05, ** P < 0.01, ***P < 0.001, ****P < 0.001). For g-j, mice were injected with MPS or bolus vaccines (GM-CSF, CpG, OVA) and dLNs were collected on at a late timepoint (day 20-21). Naïve mice were included as controls. n = 5 biologically independent animals per group, barcoded and pooled for sequencing. (g) Schematic of processing pipeline for single-cell sequencing. LNs were digested and FACS-sorted to enrich live, CD45^+^CD3^-^CD19^-^ cells for sequencing. (h) UMAP clustering of 20,858 cells across conditions. (i) UMAP clustering of cell populations from each sample condition. (j) Heatmap depicting differential gene expression across clusters used to determine cell types.

Because LN expansion is known to be mediated by myeloid interactions with LN stromal cells, we next performed single-cell RNA sequencing (scRNAseq) on the LN myeloid compartment after vaccination (Fig. 3g, Supplementary Fig. 9a) [2]. LNs were examined at a late timepoint (days 20-21) to consider mediators of long-term expansion. After removal of lymphocytes and stromal cells, we identified nine clusters from the remaining 20,858 cells analyzed (Fig 3h-j). Type-2 conventional DCs (cDC2s; c0, *Sirpa, H2-Ab1*), plasmacytoid DCs (c1, *Siglech, Bst2*), migratory DCs (c2, *Ccr7, Clu*), type-1 conventional DCs (c3, *Xcr1, Clec9a*), Langerhans cells (c4, *Cd207*), plasma cells (c5, *Ighg2b, Ighg1*), inflammatory monocytes (c6, *Csf1r, Ly6c2*), neutrophils (c7, *S100a8, S100a9*), and proliferating cDC2s (c8, Top2a, *Mki67*) were identified as cell types associated with each cluster (Fig. 3j, Supplementary Fig. 9b). Consistent with the flow cytometry analysis, scRNAseq identified broad changes in LN cell populations after immunization, with notable differences based on vaccine strength (Fig. 3i, Supplementary Fig. 9c). Compared to the PBS condition, both bolus and MPS vaccines increased DC2 proportions and decreased frequencies of migratory DCs and DC1s.

Maintenance of LN expansion was associated with increased frequencies of inflammatory monocytes and plasma cells and decreased Langerhans cells (Supplementary Fig. 9c).

### MPS vaccination alters long-term gene expression of antigen-presenting cells in LNs

Given the importance of sustained antigen presentation in maintenance of LN immune responses [19, 20], we hypothesized that vaccine antigen availability and antigen-presenting cell (APC) populations may affect LN expansion. Compared to LNs of mice given the full MPS vaccine, LNs of mice given an MPS vaccine without antigen became prominently less enlarged and contracted sooner (Fig. 4a-b).

**Figure 4.**
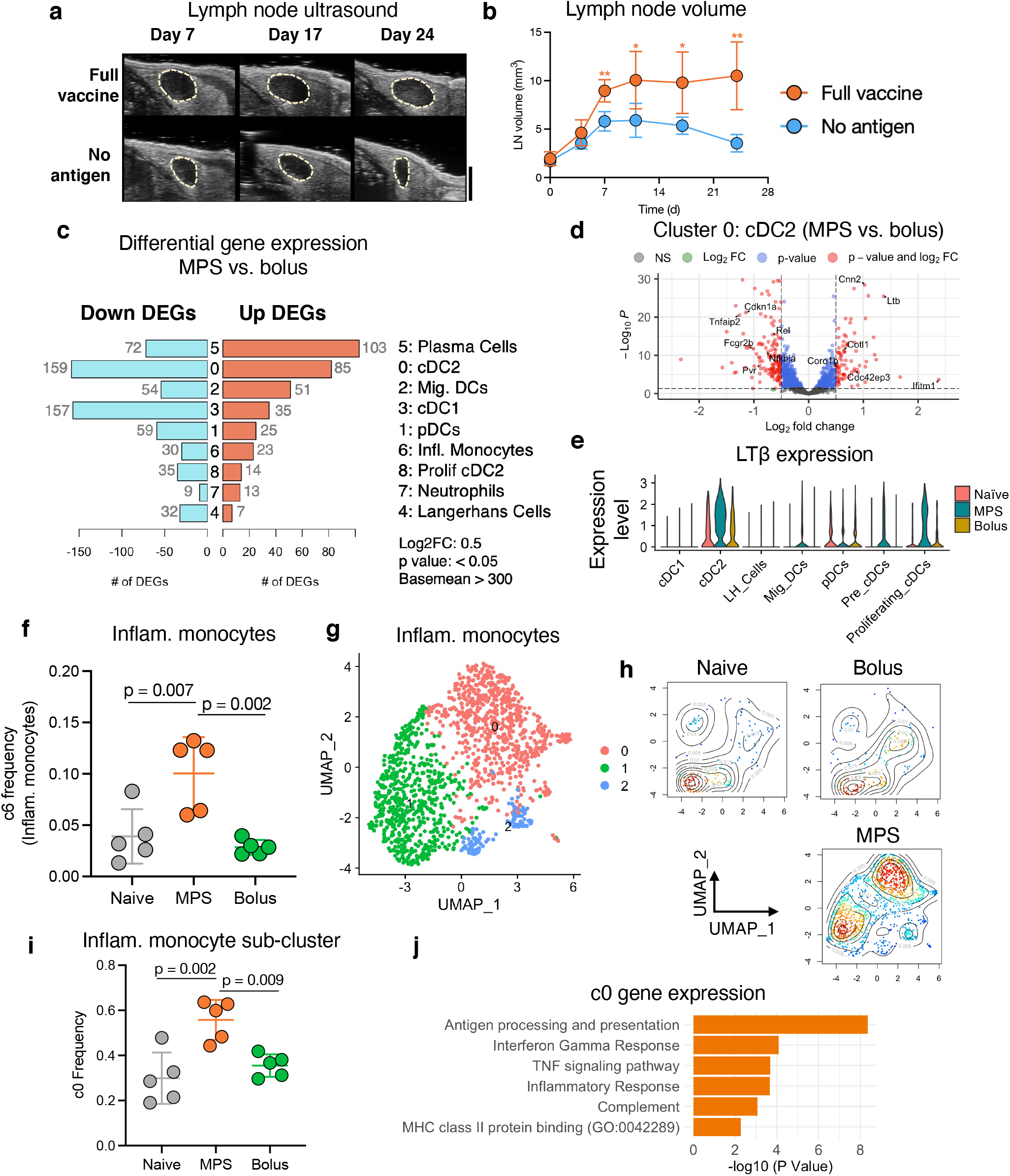
MPS vaccination elicits long-term transcriptional changes in antigen-presenting cells. (a-b) Mice were immunized on day 0 with a full MPS vaccine (containing GM-CSF, CpG, and OVA protein) or an MPS vaccine without antigen (GM-CSF and CpG only). LN volume was tracked using HFUS imaging. n = 5 biologically independent animals per group. (a) Representative HFUS images of vaccine-draining LNs. (b) Quantification of LN volume over time. Statistical analysis was performed using two-tailed t tests. (c-j) Mice were injected with MPS or bolus vaccines (GM-CSF, CpG, OVA) and dLNs were collected on at a late timepoint (day 20-21). Naïve mice were included as controls. n = 5 biologically independent animals per group, barcoded and pooled for sequencing. (c) Numbers of differentially expressed genes by cell type between the MPS and bolus conditions. (d) Volcano plot displaying differentially expressed cDC2 genes between the MPS and bolus conditions. (e) *Ltb* (lymphotoxin β) expression among DC subtypes in the different conditions. (f) Proportion of inflammatory monocytes (cluster 6 from Fig. 3h) in LNs. UMAP re-clustering of inflammatory monocyte populations among sample conditions (g), and separated by condition (h). (i) Proportion of cluster c0 among inflammatory monocytes. (j) Pathway analysis for inflammatory monocyte cluster c0. For f and i, statistical analysis was performed using analysis of variance (ANOVA) with Tukey’s post hoc test.

To identify potential mediators of this differential, antigen-dependent response, the data arising from the scRNAseq was next employed for further analysis of LN APC populations. Broadly, the cell clusters identified in scRNAseq analysis were all found to differentially express a variety of genes between the MPS and bolus vaccine conditions (Fig. 4c). The most dramatic transcriptional changes were in the LN-resident cDC2 and cDC1 compartments, moreso than migratory DCs and Langerhans cells. The cDC2s showed the greatest number of differentially expressed genes between the two vaccine strengths (Fig. 4c-d), and consensus non-negative matrix factorization (cNMF) analysis [21] identified a cDC2-specific program (CNMF_X14) enriched with MPS vaccination (Supplementary Fig. 10a-d). This program included genes involved in inflammation (*Il1r2m Cd86*), immune regulation (*Clec4a2, Sirpa, Lst1*), cell migration machinery (*Rasgef1b, Elmo1*), and smooth muscle contraction (*Ppp1r14a*) (Supplementary Fig. 10d). Furthermore, *Ltb* was strongly upregulated in cDC2s with MPS vaccination relative to the bolus condition (Fig. 4d-e, Supplementary Fig. 10d-e). MPS immunization also increased the frequency of CD19^-^ plasma cells, and directed gene expression towards more mature immunoglobulin (*Ighg1* versus *Ighm*) expression (Supplementary Fig. 11a-d).

As monocytes significantly expanded both by number and proportion following MPS vaccination (Fig. 4f, Supplementary Fig. 7p), and can present antigen to T cells in LNs [22], their gene expression was next assessed. UMAP re-clustering identified three sub-populations of inflammatory monocytes, which shifted following vaccination (Fig. 4g-h). Notably, c0 formed the predominant monocyte phenotype in LNs with sustained expansion (i.e. MPS condition), relative to naïve or bolus-vaccinated mice (Fig. 4h-i). Gene set enrichment analysis identified pathways associated with antigen processing and presentation, IFNγ response, and inflammatory signaling that differentiated monocytes in the strong and weak vaccine LNs (Fig. 4j).

### Inflammatory, antigen-presenting monocytes are associated with robust LN expansion

To confirm the impact of vaccine strength on antigen-presenting, inflammatory monocytes, LNs of mice vaccinated with the MPS vaccine (with or without antigen) were collected and further compared to LNs of mice given bolus or PBS controls (Fig. 5a).

**Figure 5.**
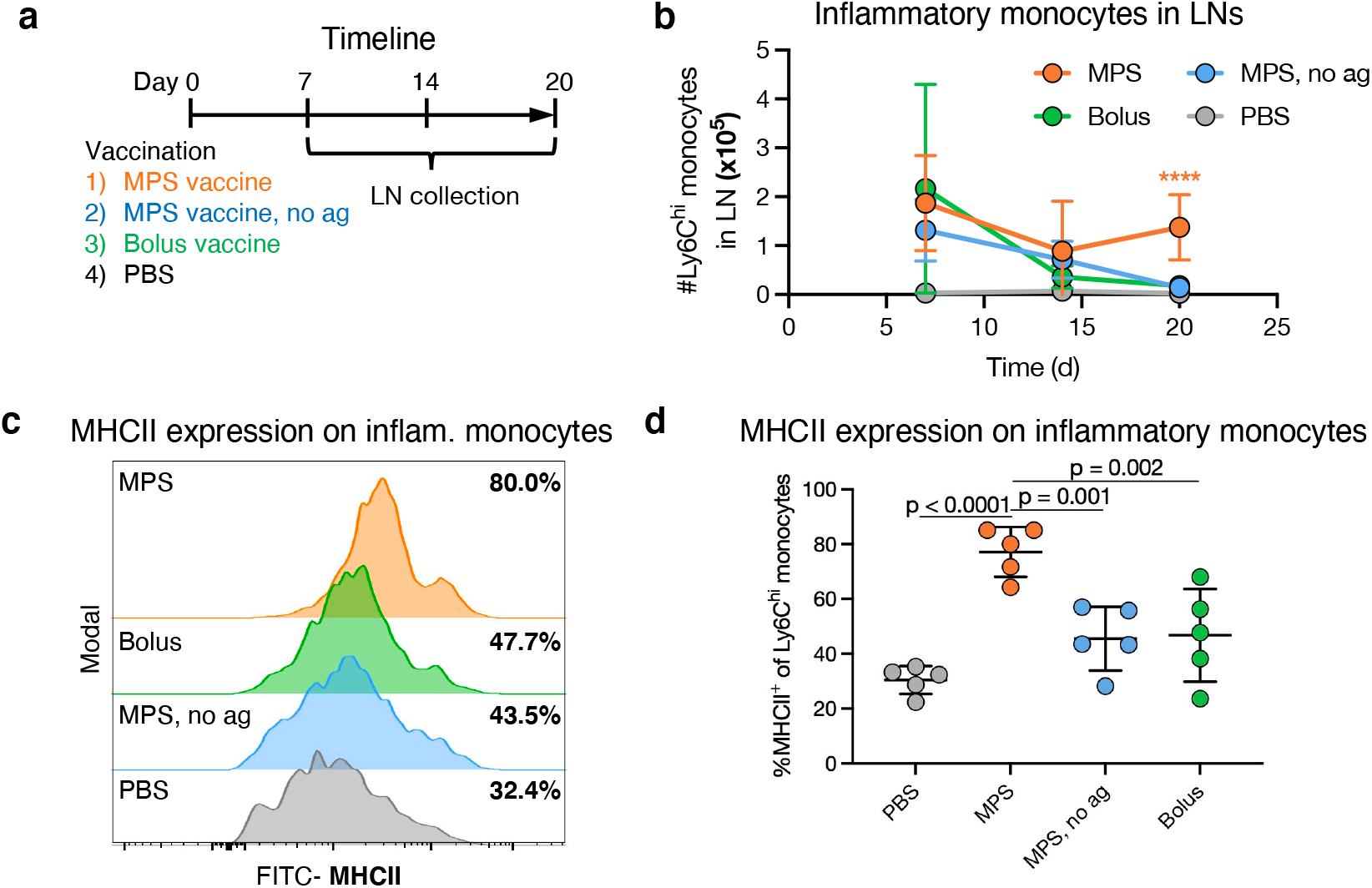
Prolonged LN expansion is associated with inflammatory monocyte engagement. Mice were treated with MPS or bolus vaccines (containing GM-CSF, CpG, OVA), MPS vaccine without antigen (GM-CSF, CpG only), or PBS, and LNs were collected on days 7, 14, and 20 for cellular analysis. n = 5 biologically independent animals per group per timepoint. (a) Experimental timeline and conditions. (b) Inflammatory monocyte number in LNs over time. (c) Representative flow cytometry histograms depicting MHCII expression on Ly6^hi^ inflammatory monocytes. Median percentage MHCII expression in each group is listed on the right. (d) MHCII expression on Ly6^hi^ inflammatory monocytes in the LN at day 20. Statistical analysis was performed using analysis of variance (ANOVA) with Tukey’s post hoc test. For b and d, means depicted; error bars, s.d. For b, statistical analysis was performed using analysis of variance (ANOVA) with Tukey’s post hoc test. For b, only differences between one group and all other groups are shown (*P < 0.05, ** P < 0.01, ***P < 0.001, ****P < 0.001).

Consistent with the scRNAseq analysis, Ly6C^hi^ inflammatory monocytes [42, 43] comprised the majority (∼60-70%) of LN monocytes in the MPS group over time, significantly higher than the PBS and bolus groups (∼40-50%) by day 20 (Supplementary Fig. 12a-b). Inflammatory monocytes were also significantly expanded by number and proportion in the LNs of MPS-vaccinated mice at day 20 compared to the PBS and bolus, and were visualized in LNs through CCR2 expression (Fig. 5b, Supplementary Fig. 12c-d) [23]. Inflammatory monocyte responses were abrogated at later timepoints when the MPS vaccine was delivered without antigen, equivalent to the PBS or bolus controls by day 20 (Fig. 5b).

Inflammatory monocytes in the MPS group displayed characteristics of antigen presentation; MHCII expression significantly increased in the MPS vaccine group, compared to all others (Fig. 5c-d). Numbers of monocyte-derived DCs (CD11c and MHCII-expressing Ly6C^hi^ monocytes) were also significantly increased in the MPS-vaccinated dLN at this time (Supplementary Fig. 13).

### Harnessing LN expansion improves vaccine outcomes

Finally, we considered whether LN expansion, and the cellular and transcriptional changes observed, might indicate functional outcomes of vaccination. In a therapeutic model of mouse melanoma, LN expansion after vaccination against a tumor-expressed antigen was not affected by tumor presence (Supplementary Fig. 14a-c). The MPS vaccine generated stronger adaptive immune responses than the bolus vaccine, leading to therapeutic benefit (Fig. 6a-c, Supplementary Fig. 14d-f, Supplementary Fig. 15a-c). Importantly, LN expansion correlated positively with antibody titers, CD8^+^ T cell responses, and antitumor efficacy of cancer vaccine formulations (Fig. 6d-f, Supplementary Fig. 16a-c). Strikingly, in a tumor-free setting, MPS vaccination also led to stronger long-term antibody (day 90) and splenic CD8^+^ T cell (day 103) responses than the bolus vaccine, and responses strongly correlated with the earlier degree of LN expansion (Supplementary Fig. 17a-f).

**Figure 6.**
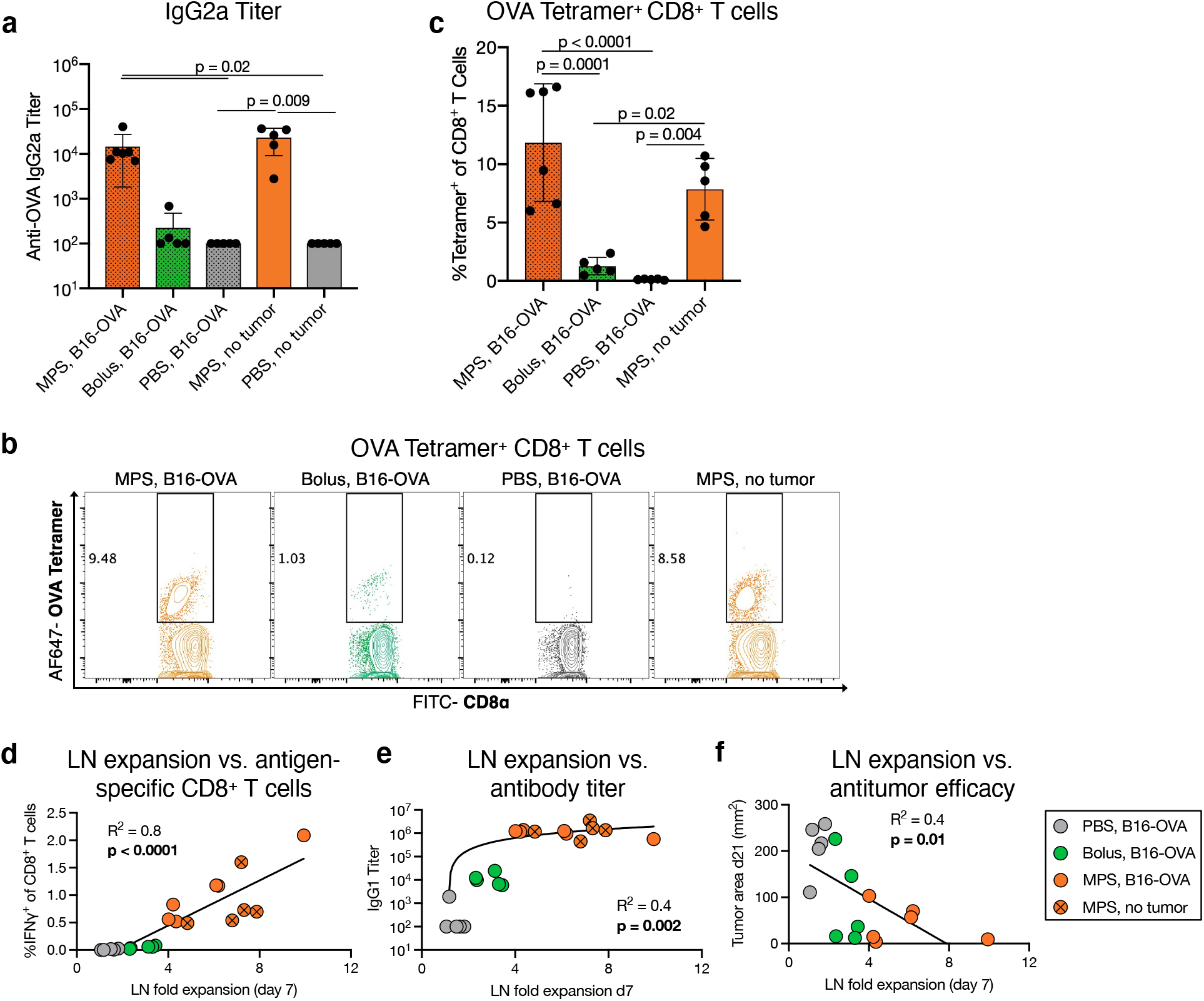
Lymph node expansion correlates with adaptive immunity and therapeutic outcomes of vaccination. Mice were inoculated with B16-OVA melanoma tumors and three days later treated with MPS or bolus vaccines containing GM-CSF, CpG, and OVA protein, and compared to PBS-injected controls. A fourth group of tumor-free mice was treated with MPS vaccines (called “MPS, no tumor”). Inguinal dLNs were imaged using HFUS at multiple timepoints, and blood was collected 8 and 21 days after vaccination to assess T cell responses and serum antibody titers, respectively. n = 5-6 biologically independent animals per group. (a) Serum anti-OVA IgG2a antibody titer, 21 days after immunization. Statistical analysis was performed using a Kruskal-Wallis test with Dunn’s post hoc test. (b-c) T cell analysis in the peripheral blood, 8 days after immunization. (b) Representative flow cytometry plots of OVA-tetramer binding to CD8^+^ T cells. (c) Proportion OVA-tetramer^+^ of CD8^+^ T cells in blood. Statistical analysis was performed using analysis of variance (ANOVA) with Tukey’s post hoc test. For a and c, means are depicted; error bars, s.d. Linear regressions are shown of LN fold expansion 7 days after vaccination versus (d) IFNγ^+^ CD8^+^ T cells after SIINFEKL restimulation, (e) anti-OVA IgG1 titers, and (f) tumor area at the latest timepoint with all mice surviving (day 21). Correlations are statistically significant. For e, the y-axis is plotted on a log scale to show the range of vaccine responses, although a linear regression was performed.

To explore if LN expansion could directly improve vaccine efficacy, the MPS vaccine without antigen (Fig. 4a-b) was employed to “jump-start” LN expansion prior to administration of a full, antigen-containing bolus vaccine (Fig. 7a). LNs of mice given the antigen-free MPS jump-start expanded over the first week and continued to increase in size after administration of the bolus vaccine, becoming significantly enlarged compared to all other groups (Fig. 7b). The jump-start plus bolus vaccine broadly improved vaccine responses compared to the traditional bolus vaccine. The proportion of OVA-tetramer^+^ CD8^+^ T cells in blood was significantly increased in this condition (Fig. 7c-d). Blood CD8^+^ T cells restimulated *ex vivo* with SIINFEKL peptide had superior cytokine production (IFNγ and TNFα) with the jump-start (Supplementary Fig. 18a-c), and the jump-start also increased the proportion of effector CD8^+^ T cells and decreased the blood CD4/CD8 T cell ratio relative to mice given the bolus vaccine alone (Supplementary Fig. 18d-e). The combination treatment improved short and long-term IgG2a antibody titers, with 10/10 (versus 6/10 with the bolus only) detectable IgG2a titers after 100 days (Fig. 7e, Supplementary Fig. 18f). B16-OVA tumor-bearing mice treated using the jump-start strategy (Supplementary Fig. 19a-b) had superior tumor control compared to the bolus vaccine, which induced only transient tumor regressions with all mice in this condition eventually succumbing to tumor burden within 50 days. In the jump-start plus bolus group, 25% of mice survived at 200 days, a significant improvement over all other groups (Fig. 7f-g).

**Figure 7.**
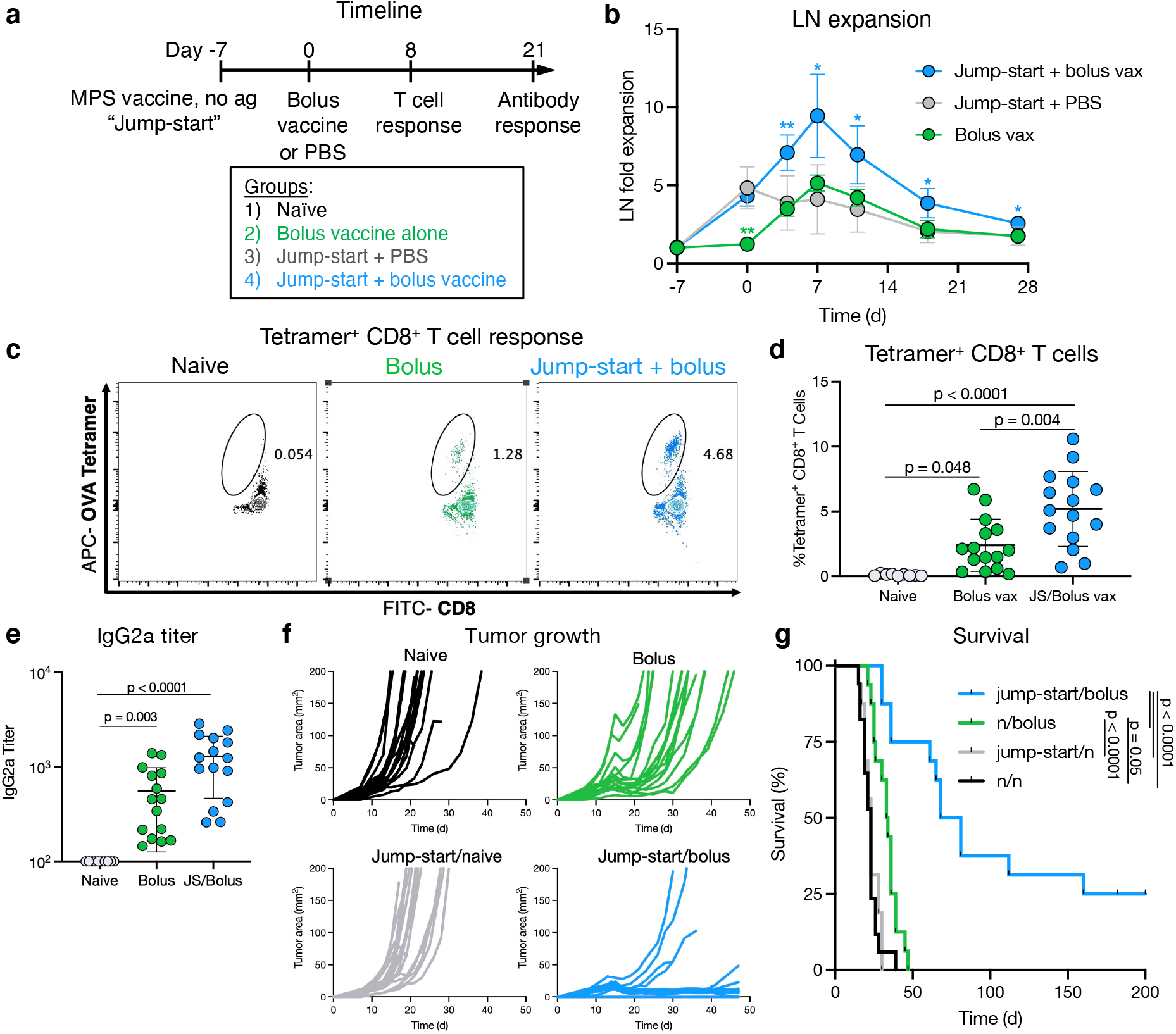
Augmenting lymph node expansion improves the magnitude and efficacy of the immune response. (a) Experimental timeline for (b-e); mice were injected with PBS or a bolus vaccine on day 0, or injected with an MPS no-antigen “jump-start” on day -7 followed by PBS or a bolus vaccine (GM-CSF, CpG, and OVA protein) on day 0. Mice were bled after 8 and 21 days for T cell analysis and serum antibody titers, respectively. (b) LN expansion measured by HFUS imaging. (c) Representative flow cytometry plots depicting CD8^+^ T cell OVA tetramer binding in cells derived from blood on day 8. (d) OVA-tetramer^+^ proportion of CD8^+^ T cells. (e) anti-OVA IgG2a titer on day 21. Statistical analysis was performed using a Kruskal-Wallis test with Dunn’s post hoc test. (f-g) Mice bearing B16-OVA tumors (inoculated day -8) were treated starting at day -7 as per studies in (a-e) with an MPS “jump-start” (MPS material, GM-CSF, and CpG without antigen) or left untreated, and then injected with a bolus vaccine (GM-CSF, CpG, and OVA) or left untreated at day 0. Tumor growth and survival were tracked. (f) Tumor growth curves. (g) Kaplan-Meier curves depicting survival. Statistical analysis was performed using a log-rank (Mantel-Cox) test, correcting for multiple comparisons. For b and d, statistical analysis was performed using analysis of variance (ANOVA) with Tukey’s post hoc test. For b, n = 5 biologically independent animals per group. For c-e, n = 10-15 biologically independent animals per group; results are combined from two independent experiments. For f and g, n = 16-17 biologically independent animals per group; results are combined from two independent experiments, the second performed in a blinded manner. For b, d, and e, means depicted; error bars, s.d. For b, only differences between one group and all other groups are shown (*P < 0.05, ** P < 0.01, ***P < 0.001, ****P < 0.001).

## Discussion

The results of this study indicate that while adjuvant and biomaterials engage early LN expansion, antigen-specific responses are key to maintenance of LN enlargement. The MPS vaccine used here as a prototypical “strong” vaccine led to a mean ∼7-fold increase in dLN volume that was maintained for weeks after immunization. This is a striking outcome compared to reported vaccine formulations (∼4-5-fold increase in mass or size, contracting from day 7-14) [2, 24–29]. In a recent study, popliteal LNs expanded ∼10-fold to vaccination with complete Freund’s adjuvant, although this response was not explored beyond 14 days [10]. The prolonged LN expansion observed here likely derives from persistence of the MPS vaccine and its sustained release of antigen and adjuvant [15]. Sustained antigen release can prolong germinal center activity in the LNs and may maximize LN expansion [19, 20, 24, 29], and it was previously shown that early explant of the MPS scaffold site impaired antibody titers [30]. By day 20, the MPS formulation led to mature Ig expression by LN plasma cells while the bolus vaccine did not, suggesting maintained germinal center responses.

The current results are, to our knowledge, the first characterization of LN spatial mechanical properties after vaccination, and reveal that vaccine draining LNs demonstrate location-specific alterations in ECM and tissue mechanical properties with immunization. Our findings are in general agreement with a previous study suggesting LN softening during podoplanin blockade [8]. The consistent cell density and spatial arrangement found here in LNs during enlargement suggests that LNs expand coordinately with the degree of internal cellular expansion, and are not due to changes in cell density. The relaxation of individual FRCs, increasing gap size during LN expansion, or altered ECM distribution could instead be responsible for this phenomenon [2, 6, 10, 31]. Previously, parallel plate compression of explanted LNs recorded a modest increase in Young’s modulus following vaccination, potentially due to this technique capturing internal proliferation and pressure, or fibrosis of the LN capsule [10].

Inflammatory monocytes are strongly implicated as key mediators of the strength of the MPS vaccine response. During LN expansion following MPS vaccination, immune and stromal cell numbers increased in draining LNs, while weaker vaccination (i.e. bolus) led to minimal cellular expansion. Immune cell numbers peaked earlier than stromal cells (day 4-7 versus 11), consistent with prior vaccine work [2, 7, 9]. Strikingly, Ly6C^hi^ inflammatory monocytes rapidly increased in LNs following ‘strong’ vaccination and remained elevated throughout LN expansion. These monocytes upregulated antigen-presentation related genes in the MPS relative to the bolus vaccine group, supporting a role in antigen presentation [22]. Monocytes can also affect the quality of T cell response (e.g. effector phenotype) through cytokine secretion and T cell co-localization, and mobilize in response to type I adjuvants including CpG [32].

The upregulation of genes associated with LN development and maintenance (*Ltb*) in cDC2s with MPS vaccination suggest a role of cDC2s in prolonging LN expansion. Ltb encodes lymphotoxin β, with known involvement in lymphogenesis and expansion [9]. Although DCs were previously associated with driving LN expansion, the exact subtype was not defined, relying on a mouse model with broad CD11c^+^ cell modification [2]. Here, cDC2s were the main differentiator between the ‘strong’ and ‘weak’ vaccine conditions in terms of differential gene expression, and upregulated *Ltb*. Because the most dramatic changes in gene expression occurred with LN-resident APC populations (cDC1s, cDC1s, plasma cells), these results suggest that the MPS vaccine strongly and specifically affects the LN environment to produce strong protective immunity.

Importantly, LN expansion correlated with therapeutic vaccine outcomes, and expansion of LNs prior to bolus vaccination significantly improved efficacy. Previously, in a model of E7 peptide vaccination in C3 HPV-E7 tumor-bearing mice, vaccine response was reported to correlate with LN expansion (assessed by MRI) [24]. Accordingly, minimally invasive LN imaging technologies such as HFUS or MRI may indicate the strength of vaccine immune responses in mouse models. In humans, certain vaccines such as those against SARS-CoV-2 can lead to lymphadenopathy in a subset of patients [33, 34]. It remains to be shown whether this expansion indicates initiation of a successful immune response. The jump-start vaccine concept developed here comprises generalized, off-the-shelf components and could potentially be applied to broadly improve subsequent antigen-specific therapy.

## Methods

### MPS vaccine fabrication

MPS vaccines were prepared as previously described [15, 30]. Briefly, 4g of Pluronic P123 surfactant (average M_n_ ∼5800, Sigma) was dissolved in 150g of 1.6M HCl, mixed with 8.6g tetraethyl orthosilicate (TEOS, Sigma) at 40°C for 20h, and stirred at 100°C for another 24h to fabricate MPS. Pluronic P123 was cleared in 1% ethanol in HCl at 80°C for 18h, and MPS was filtered and dried at 65°C. To prepare MPS vaccines, MPS was suspended in pH 7.4, sterile DPBS, and 2mg was incubated each with OVA (200μg/vaccine, unless otherwise indicated in experiments using MPS vaccines without antigen) or CpG (100μg/vaccine) at room temperature for ∼7h before freezing and overnight lyophilization. The next day, a separate 1mg aliquot of MPS was suspended in DPBS and loaded with GM-CSF (1μg/vaccine) for 1h at 37°C while shaking. The three MPS populations (loaded with antigen, CpG, and OVA) were combined in 150μL of sterile DI H_2_O. Each MPS vaccine mixture was injected through an 18G needle into the left flank of the mouse. GM-CSF was purchased from PeproTech (Rocky Hill, NJ). CpG-ODN 1826 5′-TCCATGACGTTCCTGACGTT-3′ was synthesized by Integrated DNA Technologies (Chicago, IL). OVA was purchased from InvivoGen (InvivoGen vac-pova, San Diego, CA). Vaccine components (MPS rods, CpG, GM-CSF, and OVA) were endotoxin tested (<0.01 EU/vax).

### Bolus vaccines

OVA (200ug/vaccine), CpG (100μg/vaccine), and GM-CSF (1μg/vaccine) were mixed in pH 7.4, sterile DPBS to obtain a total volume of 150μL per dose.

### Ultrasound imaging

Mice were anesthetized prior to imaging. Hair on the left and right abdomen was shaved and fully removed using Nair hair removal lotion. Mice were placed on a heated stage (37°C) and their limbs were gently secured using tape (3M Transpore). Signa creme electrode cream (Parker Laboratories) was applied on the paws to detect respiration. The stage was rotated about the long axis of the animal (∼30°) to expose the inguinal region, and Aquasonic ultrasound transmission gel (Parker Laboratories) was applied. Inguinal LNs were imaged using a Vevo 3100 scanner and 50 MHz transducer on a Vevo 3100 Preclinical Imaging System (Visualsonics).

Respiration gating was enabled to avoid respiratory artefacts and LNs were scanned using a step size of 0.1mm. LN volumes were quantified using Vevo LAB software (Visualsonics). This procedure was also applied to image the vaccine site. To account for the effect of circadian rhythms in lymphocyte trafficking [35, 36], LNs were imaged at the same time of day (+/-1h) for the duration of each experiment.

### Cellular analysis of lymph nodes

At days 4, 7, 11, 17, 51 or 7, 14, 20 post-vaccination, mice were euthanized and vaccine draining and non-draining LNs were explanted and digested in RPMI-1640 (Corning) containing 0.8mg/mL Dispase II, 0.2mg/mL Collagenase P, and 0.1mg/mL DNase I (all Roche, procured from Sigma) until no visible LN pieces remained, following established protocols [37, 38]. Cell suspensions were strained through a 70μm filter, counted using a Countess II FL (ThermoFisher), and stained using standard flow cytometry protocols using antibodies in Table 1. Viability was assessed using a Zombie Yellow stain (BioLegend) and samples were run on an Aurora Spectral Analyzer (Cytek) or LSRII flow cytometer (BD Biosciences). Flow cytometry gating strategies for immune cell populations are indicated in Supplementary Fig. 8.

**Table 1.**
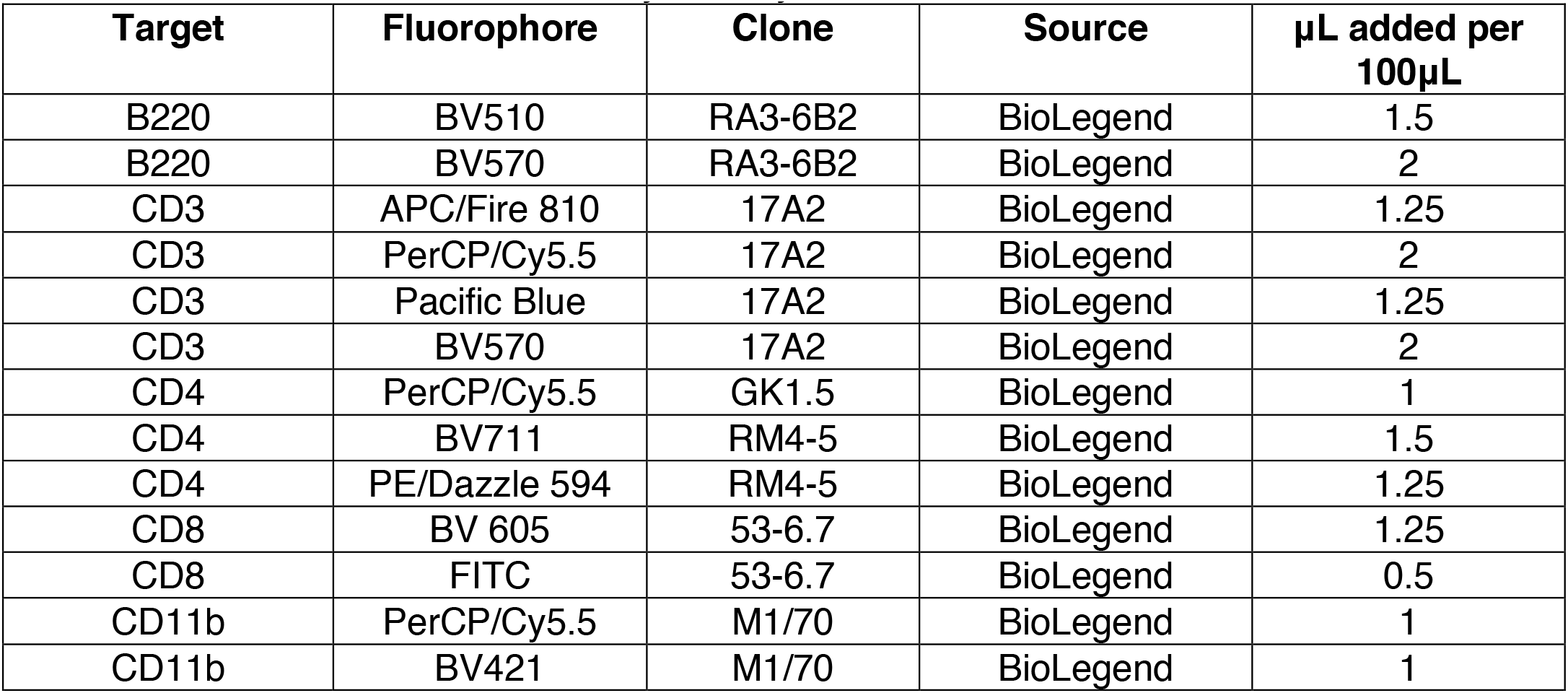
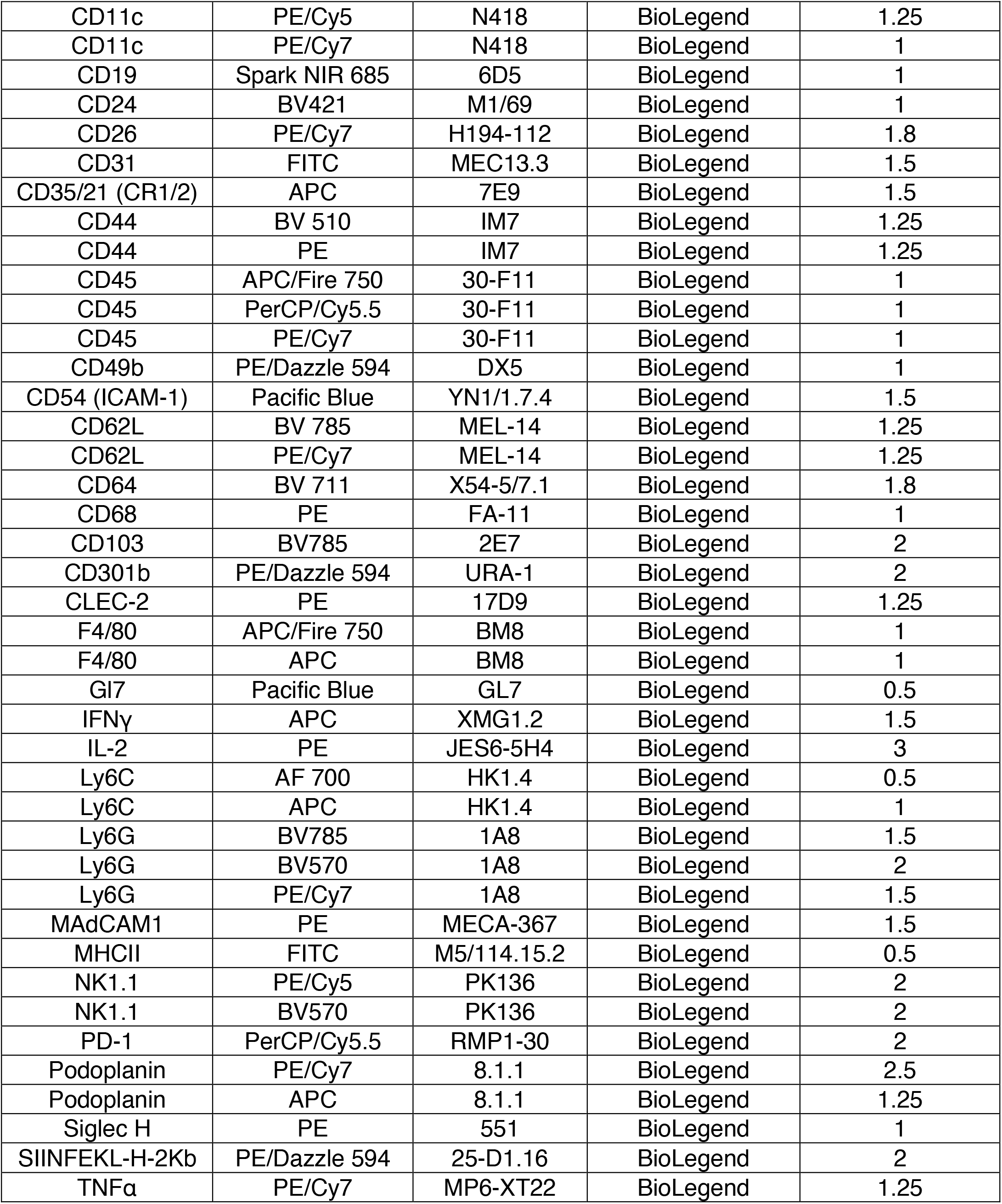
Antibodies utilized in flow cytometry

### Peripheral blood analysis

Blood was collected from mice retro-orbitally using heparinized capillary tubes (Fisherbrand) and stored in heparinized collection tubes (BD Biosciences) on ice. Red blood cells were lysed using ACK Lysing Buffer (Quality Biological). To detect OVA-specific CD8^+^ T cells, samples were incubated with 7μg/mL allophycocyanin (APC)-conjugated H-2Kb-SIINFEKL tetramer (National Institutes of Health tetramer core facility) at 37°C. For cytokine analysis, cells were instead incubated with 2μg/mL OVA257–264 SIINFEKL OVA CD8 peptide for 1.5h at 37°C. GolgiStop (BD Biosciences) was then added and samples incubated for another 4h at 37°C. Samples were stained with anti-mouse CD3, CD8, and CD4 antibodies, and treated with a Cytofix/Cytoperm kit (BD Biosciences) per manufacturer protocol.

Permeabilized samples were stained for IFNγ, IL-2, and TNFα and analyzed by standard flow cytometry protocols. For general cellular analysis, blood samples were stained with antibodies from Table 1.

### Antibody titers

Blood was collected from mice and allowed to coagulate for 30 minutes at room temperature. Tubes were centrifuged at 2200xg for 10min and serum collected and transferred into low-binding tubes for storage at -80°C. Anti-OVA antibody titers were quantified using ELISA. Briefly, high-binding plates (Costar 2592, Cole Palmer) were coated at 4°C overnight with 10μg/mL OVA in DPBS with gentle rocking. Serum samples were diluted (1:10^2^-1:10^8^) in 1x ELISA diluent in pH 7.4 DPBS and added to the washed plate for 2h at room temperature. Plates were washed and biotinylated anti-mouse IgG1 and IgG2a antibodies (BD Biosciences, each diluted 1:100 in 1x ELISA diluent) were added for 2h at room temperature. Plates were washed and streptavidin-HRP (BD Biosciences, diluted 1:1000 in 1x ELISA diluent) was added for 15 minutes at room temperature. Plates were again washed and TMB substrate solution (BioLegend) was added for 15 minutes at room temperature prior to addition of stop solution (2M hydrochloric acid). Absorbance was read at λ = 450nm, subtracting background λ = 540nm. Anti-OVA titer was defined as the lowest serum dilution with OD = 0.2.

### Immunohistochemistry of lymph nodes

Mice were euthanized and inguinal LNs were harvested into 0.25mL PBS on ice and cleaned of surrounding tissue. LNs were fixed in 1% paraformaldehyde in PBS for 1 hour at 4°C, rinsed 3x with PBS, and transferred into 30% sucrose in PBS overnight at 4°C. LNs were subsequently transferred to a 1:1 solution of 30% sucrose in PBS and Tissue-Tek O.C.T. compound (OCT, VWR) for 45min at room temperature. LNs were blotted on a Kimwipe, added to OCT-containing cryomolds on dry ice, and stored at -20°C. 15μm sections were prepared using a cryostat (Leica). For staining, samples were placed in PBS for 2 minutes to dissolve OCT and blocked in 3% normal goat serum and 1% BSA in PBS at room temperature. When biotin-containing stains were used, samples were also blocked in 0.01% avidin and 0.001% biotin in subsequent steps, both for 15 minutes at room temperature.

Primary antibody staining solution (Table 2) was added overnight at 4°C. The next day, secondary and nuclear stains including streptavidin-AF594 conjugate (Invitrogen, 1:100), AlexaFluor 647 goat ant-rabbit IgG (Invitrogen, 1:100) and Hoechst 33342 (Thermo Scientific, 1:500) were added for 2 hours at room temperature. Samples were mounted using ProLong Gold antifade reagent (Invitrogen) and imaged using a Zeiss 710 Confocal system. Images were analyzed using ImageJ software and custom code in MATLAB.

**Table 2.** Antibodies utilized in lymph node immunohistochemistry

| Target | Conjugate | Clone/Catalog # | Source | Dilution |
| --- | --- | --- | --- | --- |
| B220 | AF594 | RA3-6B2 | BioLegend | 1:50 |
| CCR2 | n/a | EPR20844 | Abcam | 1:50 |
| CD3 | AF647 | 17A2 | BioLegend | 1:100 |
| CD11b | AF488 | M1/70 | BioLegend | 1:100 |
| Collagen I | n/a | AB765P | Millipore Sigma | 1:40 |
| Collagen VI | AF488 | ER-TR7 | Santa Cruz<br>Biotechnology | 1:50 |
| F-actin<br>(phalloidin) | AF488 | A12379 | Thermo Fisher<br>Scientific | 1:100 |
| Hyaluronic acid | Biotin | 385911 | Millipore Sigma | 1:50 |
| Ly6C | Biotin | ER-MP20 | Abcam | 1:200 |

### Nanoindentation of lymph nodes

Mice were injected with an MPS vaccine or PBS control on day 0, and euthanized on days 4, 7, or 11. Inguinal vaccine-draining LNs were harvested, embedded in 4% agarose in cryomolds, sectioned into 500μm thick sections using a vibratome (Leica), and collected on glass slides [39–41]. Sections were kept hydrated in PBS and stored on ice for the duration of the procedure. Mechanical testing was performed using a G200 nanoindenter (Keysight Technologies) with a 400μm spherical tip [42]. Sample points were defined in the center and periphery of each section, and PBS was periodically added to maintain hydration. For each LN, outlier data points were identified through ROUT test (Q = 1%) in GraphPad Prism V9 software and excluded from analysis. To generate mechanical maps of LN tissue, LN slices were first photographed and imported into Adobe Illustrator v22.1 software to trace the perimeter. The LN edge was identified visually using the lens on the G200 nanoindenter; sample points spanning the LN were then selected, and x-y coordinates were collected and point location recorded in Illustrator. Indentation points were spaced to avoid tip overlap (minimum spacing between nearest points ∼240 +/-46 μm) and the agarose mold (peripheral points selected ∼150μm from the LN edge). Visual maps were prepared by coloring the space closest to each sample point on a gradient corresponding to its stiffness measurement.

### Jump-start experiments

Mice were injected with MPS loaded with CpG and GM-CSF but no antigen. Seven days later, mice were administered a bolus vaccine or a PBS control. This vaccine strategy was applied to naïve mice or mice bearing B16-OVA tumors (injected one day prior to jump-start). Peripheral blood was collected 8 and 21 days after vaccination to assess antigen-specific CD8^+^ T cell and antibody responses, respectively, or tumor growth and animal survival were monitored.

### Tumor studies

B16-OVA cells were cultured in DMEM (Gibco) containing 10% FBS (Sigma) and 0.4mg/mL G418 selection antibiotic (Gibco). Cells were collected, counted using a hemocytometer, and resuspended in cold PBS. 2.5×10^5^ (MPS tumor experiments) or 1.25×10^5^ (jump-start tumor experiments) B16-OVA cells were injected subcutaneously using a 25G needle into the upper flank of mice. Beginning at day 7, tumor sizes were measured externally using calipers, and area calculated A = (π/4) * length * width. Mice were tracked and euthanized per IACUC protocol according to a cumulative score incorporating body condition, weight loss, and tumor size (tumors ≥ 17mm in any two dimensions).

### Single-cell sequencing setup and sorting

Mice were treated with an MPS or bolus vaccine (containing GM-CSF, CpG, and OVA) and draining LNs were collected on day 7 (MPS) or 20 (MPS and bolus) and compared to naïve controls. LNs were digested in RPMI-1640 (Corning) containing 0.8mg/mL Dispase II, 0.2mg/mL Collagenase P, and 0.1mg/mL DNase I (all Roche, procured from Sigma) until no visible LN pieces remained and passed through a 70μm filter. Samples were incubated in TruStain FcX (Fc block, 1:200, BioLegend) prior to incubation with staining antibodies (PE/Cy7 anti-mouse CD45, BV711 anti-mouse CD3, BV421 anti-mouse CD19, FITC anti-mouse CD31, and APC anti-mouse podoplanin, all BioLegend) and one Hashtag antibody per replicate using BioLegend TotalSeq-B03 (catalog numbers 155831, 155833, 155835, 155837, 155839; barcode sequences ACCCACCAGTAAGAC (-B0301), GGTCGAGAGCATTCA (-B0302), CTTGCCGCATGTCAT (-B0303), AAAGCATTCTTCACG (-B0304), CTTTGTCTTTGTGAG (-B0305), all diluted 1:100 in FACS buffer). Cell viability was assessed through eBioscience Fixable Viability Dye eFluor 780 (ThermoFisher). Cells were gently washed and resuspended in PBS + 2% FBS prior to sorting. Myeloid (viable, CD45^+^CD3^-^CD19^-^) and stromal (viable, CD45^-^ FSC-A^hi^) cells were enriched using a BD FACS Aria cell sorter into RPMI + 30% FBS on ice. Cells were counted and resuspended at an adequate concentration for loading.

Samples were prepared using a Chromium Single Cell 3’ Reagent Kit (10x Genomics) and run on a NovaSeq S4-200 cell (Illumina).

### Single-cell sequencing analysis

Single-cell RNA-seq data for each library from each cell type was processed with CellRanger *count* (10x Genomics) using a custom reference gene annotation based on mouse reference genome GRCm38/mm10 and GENCODE gene models. UMI count tables were read in R with the Seurat [64] package and counts were normalized to (CPM/100+1) and log transformed. UMI counts of barcode antibodies to label individual replicates were first normalized with centered log ratio (CLR) transformation per cell (Seurat, *NormalizeData*), followed by antibody assignment where the top 95% quantile hashtags were considered a positive signal (*HTODemux*). Individual libraries were merged into a Seurat object. Cells with less than 300 measured genes or over 5% mitochondrial counts were removed. Only cells labeled as “singlets” based on HTO counts were included. The top 2,000 most variable genes were selected via variance stabilizing transformation (vst) (*FindVariableFeatures*) and their expression was scaled (*ScaleData*). Principal component analysis was then performed in this gene space (*RunPCA*). Clustering was carried out based on the shared nearest neighbor between cells (*FindNeighbors*) and graph-based clustering (*FindClusters*). For graph-based clustering and generation of a UMAP reduction (*RunUMAP*) the same number of principal components was used as input [64]. 51,618 cells were used to identify clusters. Residual T cell and B cell clusters were removed from the analysis, since they were initially negatively selected prior to sequencing. For the remaining 44,897 cell objects, 20 PCs and a resolution of 0.1 were used to identify clusters. Clusters were annotated as DC2 cells (*Sirpa, H2-Ab1*), NK cells (*Nkg7, Kirk1*), migratory DCs (*Ccr7, Cd207*), plasmacytoid DCs (*Bst2, Siglech*), DC1 cells (*Xcr1*), monocyte/macrophages (*Csf1r*), fibroblastic reticular cells (*Pdpn, Ccl19*), plasma cells (*Ighg2b*), neutrophils (*Cx3cr1*), blood endothelial cells (*Pecam1, Aqp1*), and lymphatic endothelial cells (*Lyve1, Pdpn*). Data from individual populations (e.g monocyte/macrophages, DCs) were reprocessed in Seurat as described above.

Markers for each cluster were detected using the FindAllMarkers function in Seurat with default parameters. The average expression of markers was calculated for each cluster using the *AverageExpression* function in Seurat, and per-gene z-scores were calculated for visualization with the pheatmap package. Density plots of cells were generated using the UMAP coordinates of cells from each condition using the LSD R package (https://cran.r-project.org/web/packages/LSD/index.html).

Cluster population frequency changes for each cluster were evaluated with Dunn’s test with p-values adjusted via Benjamini-Hochberg following a positive Kruskal– Wallis test. Marker genes for each cluster were identified if they were expressed in at least 10% of cells in the cluster with log(0.25) fold or higher compared to all other clusters (*FindAllMarkers*).

NMF programs were identified using cNMF (package https://github.com/dylkot/cNMF#consensus-non-negative-matrix-factorization-cnmf-v14; [21]) using raw counts as input. cNMF was run with 20 iterations, 2000 genes, a local density threshold of 2, and all other options were left as default. Optimal K was chosen from between 4 to 20 as the K with a local maximum stability.

## Acknowledgements

We thank Chiara Gerhardinger and Zachary Niziolek for assistance with scRNAseq and FACS, Jack Alvarenga for support with nanoindentation, and Dr. Kwasi Adu-Berchie, Dr. David Zhang, and Dr. Aileen Li for valuable scientific discussions. This work was supported by the National Institutes of Health/National Cancer Institute (R01 CA223255). A.J.N. recognizes a Graduate Research Fellowship from the National Science Foundation. B.R.F. recognizes support from the NIH/NIA (K99 AG065495).

A.E.-A. received funding for this work from the European Union’s Horizon 2020 research and innovation programme through a Marie Sklodowska-Curie grant agreement no. 798504.

