## Supplemental data for "Lymph node expansion predicts magnitude of vaccine immune response"

**Supplemental figures for "Lymph node expansion predicts magnitude of vaccine immune response"**

**Supplementary Figure 1.** MPS vaccination induces robust, prolonged LN expansion.
**Supplementary Figure 2.** F-actin distribution in LNs.
**Supplementary Figure 3.** Mapping LN viscoelasticity.
**Supplementary Figure 4.** LN mechanical properties vary with location.
**Supplementary Figure 5.** The LN mechanical response to immunization depends on tissue location.
**Supplementary Figure 6.** LN density after immunization.
**Supplementary Figure 7.** Immune and stromal cell populations in LNs after vaccination.
**Supplementary Figure 8.** Representative gating strategy to identify immune cell subsets.
**Supplementary Figure 9.** scRNAseq gating, cluster identification, and population frequencies.
**Supplementary Figure 10.** cDC2s are transcriptionally altered after MPS vaccination.
**Supplementary Figure 11.** Plasma cells are enriched and express mature Ig after MPS vaccination.
**Supplementary Figure 12.** MPS vaccination expands inflammatory monocytes.
**Supplementary Figure 13.** Monocyte-derived DCs in LNs.
**Supplementary Figure 14.** Therapeutic study to assess correlations of LN expansion with vaccine efficacy.
**Supplementary Figure 15.** Vaccine therapeutic efficacy.
**Supplementary Figure 16.** The adaptive, antitumor vaccine response correlates with LN expansion.
**Supplementary Figure 17.** Long-lived vaccine responses correlate with LN expansion.
**Supplementary Figure 18.** Antigen-free MPS "jump-start" strategy boosts bolus vaccine response.
**Supplementary Figure 19.** Therapeutic "jump-start" experiment layout.

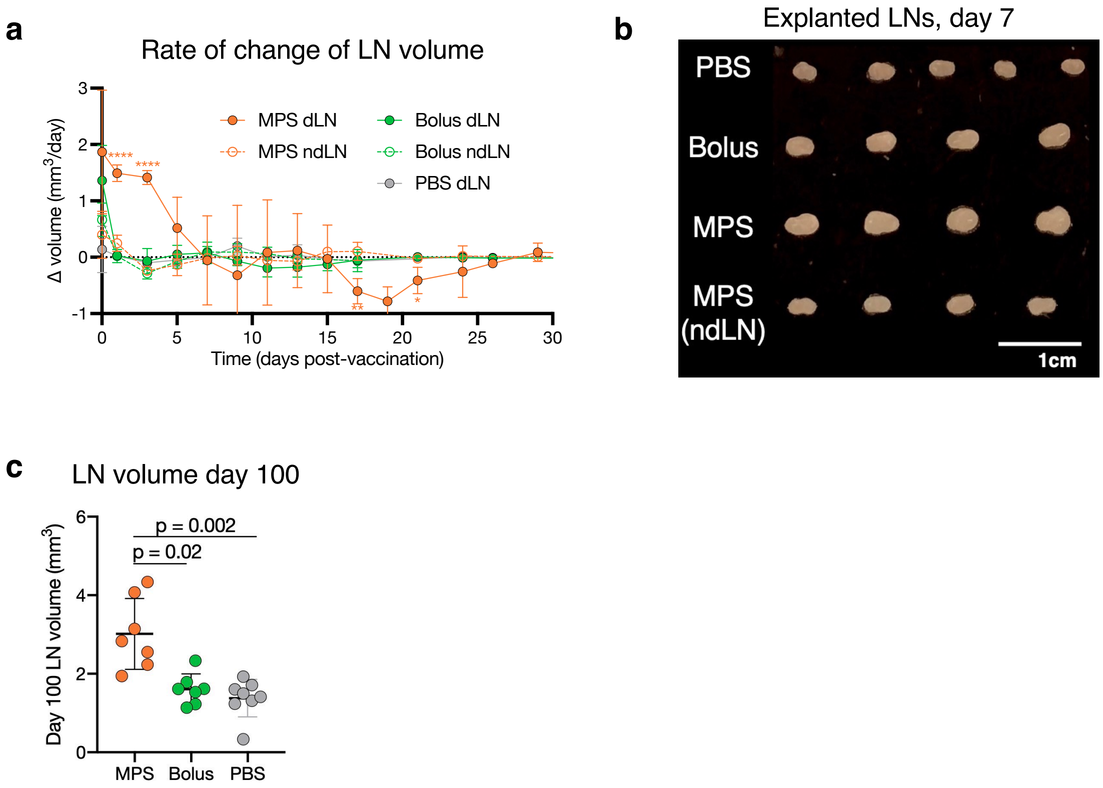


**Supplementary Figure 1. MPS vaccination induces robust, prolonged LN expansion.** Mice were immunized with MPS or bolus vaccines delivering GM-CSF, CpG, and OVA protein, and compared to PBS-injected controls. Vaccine-draining and non-draining LNs were longitudinally imaged using high frequency ultrasound. n = 7-8 biologically independent animals per group, imaged longitudinally in two cohorts. (a) Rate of change of LN volume over time. Statistical analysis was performed using analysis of variance (ANOVA) with Tukey's post hoc test. Only differences between one group and all other groups are shown (*P < 0.05, ** P < 0.01, ***P < 0.001, ****P < 0.001). (b) Photograph of LNs explanted 7 days after vaccination. Each LN was derived from a unique mouse. (c) LN volumes 100 days after immunization. Statistical analysis was performed using a Kruskal-Wallis test with Dunn's post hoc test. For a and c, means depicted; error bars, s.d.

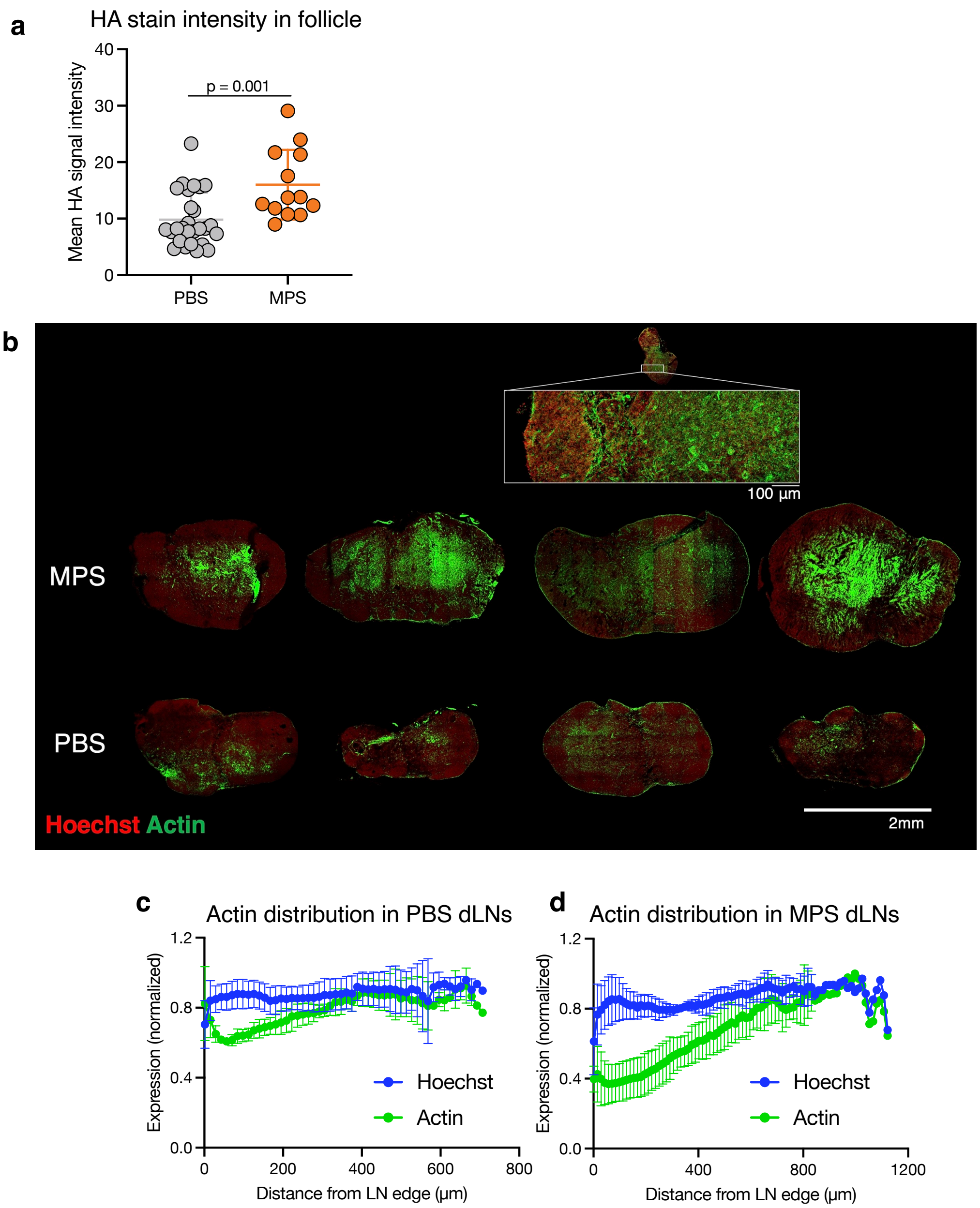


**Supplementary Figure 2. F-actin distribution in LNs.** Mice were treated with MPS vaccines (GM-CSF, CpG, OVA) or PBS and LNs were harvested after 7 days. (a) ImageJ quantification of hyaluronic acid stain intensity in follicles from PBS or MPS draining LNs on day 20. Statistical analysis was performed using a Mann-Whitney test. (b) IHC of LNs on day 7 stained for F-actin; one MPS and PBS LN from this image were selected for Figure 2c. n = 4 biologically independent animals per group. Distribution of nuclear (Hoechst) and F-actin stains across PBS (c) and MPS (d) draining LNs (from periphery to center), quantified through a custom MATLAB code, on day 7. For a and c-d, means are depicted; error bars, s.d. n = 3-4 biologically independent animals per group.

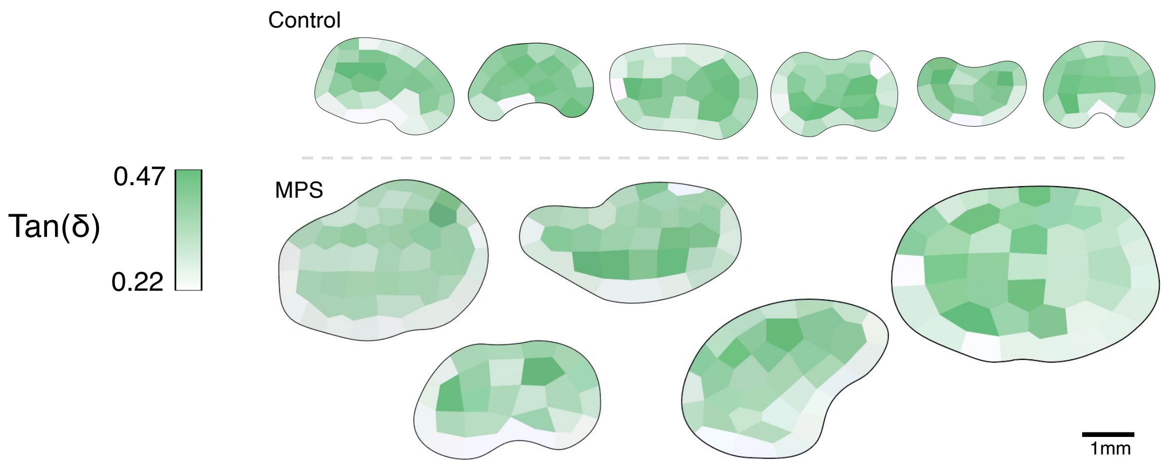


**Supplementary Figure 3. Mapping LN viscoelasticity.** Mice were treated with MPS vaccines (GM-CSF, CpG, OVA) or PBS and LNs were harvested after 7 days. Heatmaps depicting tan(δ) across individual LNs, scaled low-high within each LN (mean low value = 0.22; mean high value = 0.47). Scale bar = 1mm.

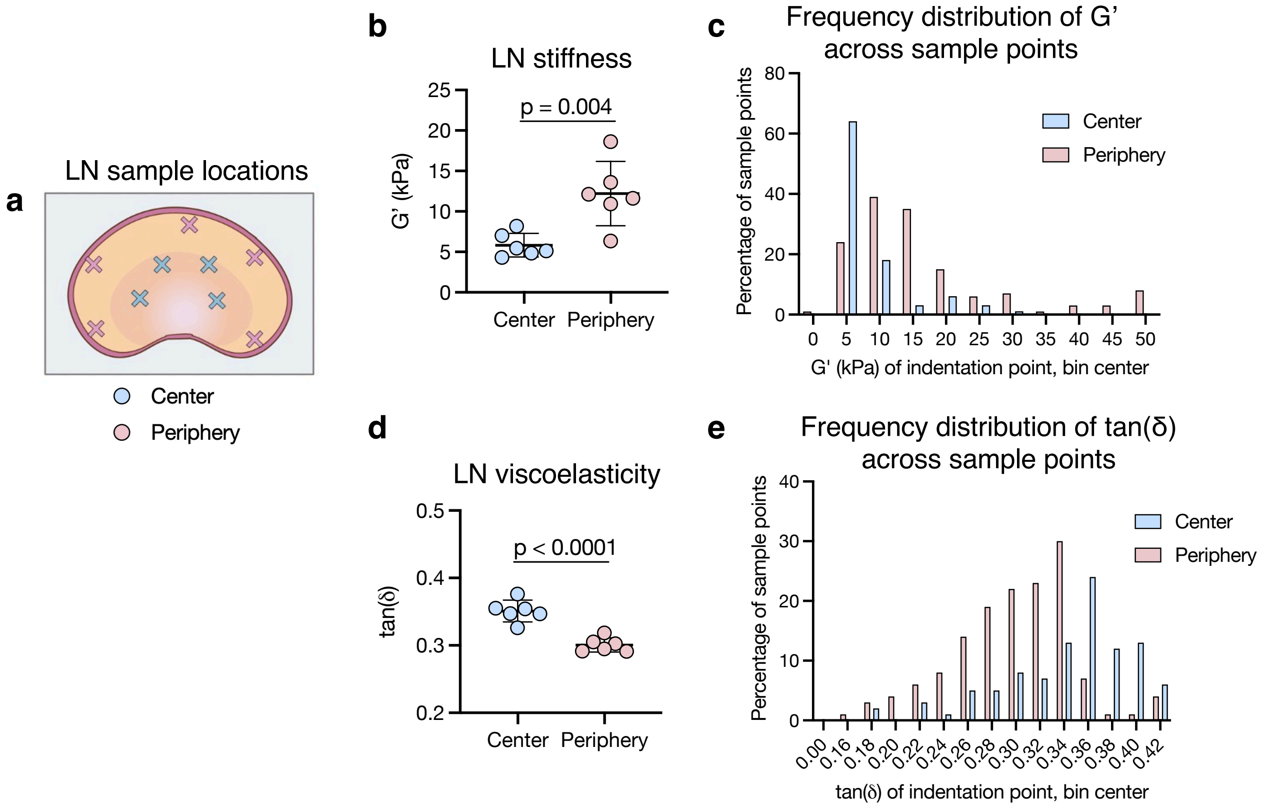


**Supplementary Figure 4. LN mechanical properties vary with location.** Inguinal LNs were collected from naïve mice for nanoindentation, and kept hydrated throughout the procedure. (a) Schematic depicting example sample locations chosen on the center or periphery. (b) Mean G' of sample points collected in the center or periphery across LNs. Each data point represents a unique LN. (c) Frequency distribution of G' of individual samples points taken across control (naïve or PBS) LNs from multiple experiments. (d) Mean tan(δ) of sample points collected in the center or periphery across LNs. Each data point represents a unique LN. (e) Frequency distribution of tan(δ) of individual samples points taken across control (naïve or PBS) LNs from multiple experiments. n = 6 independent LNs; means are depicted; error bars, s.d. For b and d, statistical analysis was performed using a two-tailed t test.

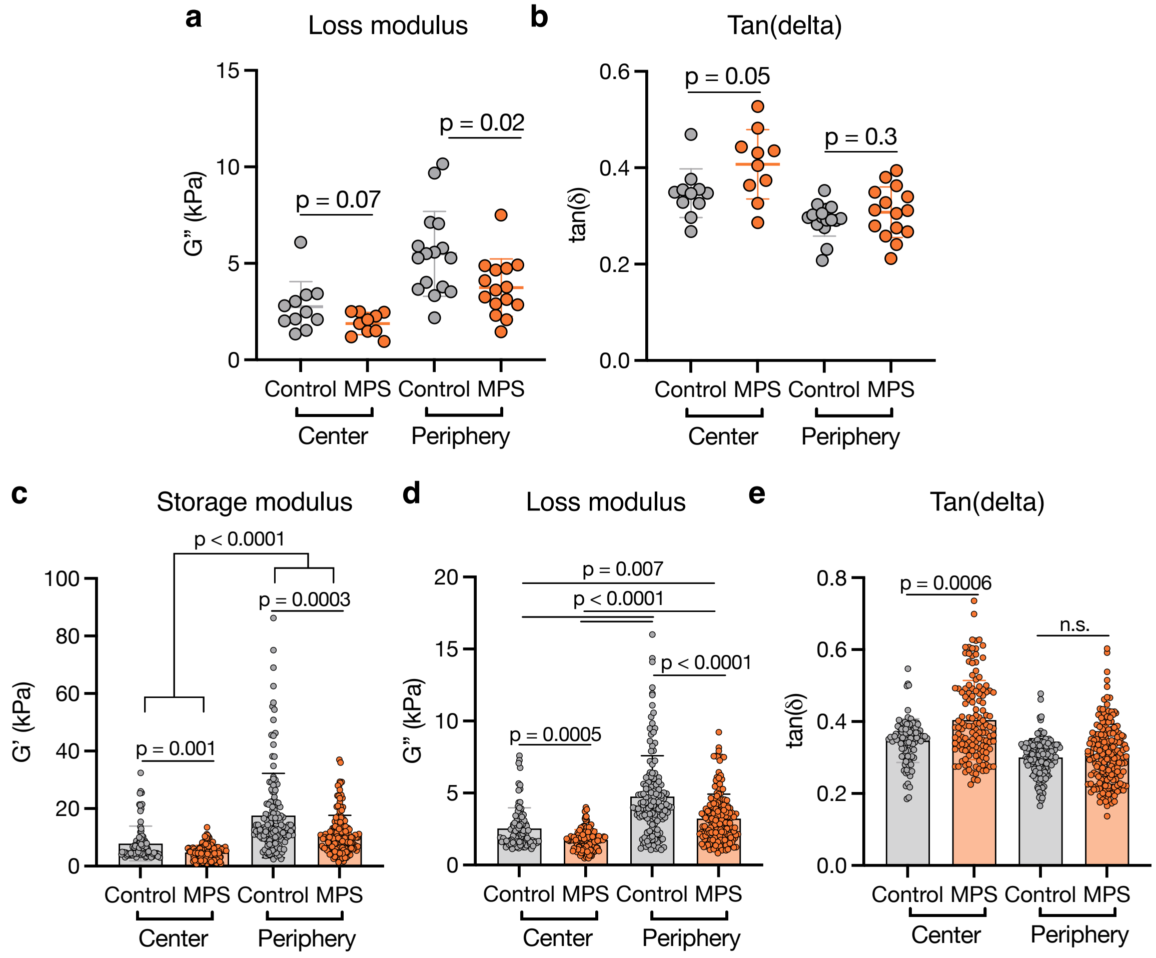


**Supplementary Figure 5. The LN mechanical response to immunization depends on tissue location.** Mice were immunized with MPS vaccines (containing GM-CSF, CpG, OVA) and dLNs were collected on day 7 for nanoindentation, and compared to PBS-injected mice or naïve controls. Mean (a) G'', and (b) tan(δ) of sample points across each LN. For a-b, each data point represents a unique LN/mouse; n = 10-16 biologically independent animals per group. Individual sample points plotted for (c) G', (d) G'', and (e) tan(δ) collected from multiple LNs. For c-e, each data point represents a single nanoindentation location, with multiple across individual LNs. For a-e, results are combined from three independent experiments; means depicted; error bars, s.d. Statistical analysis was performed using a Mann-Whitney test (a-b (center) and e), two-tailed t test (a-b (periphery)), and Kruskal-Wallis test with Dunn's post hoc test (c-d).

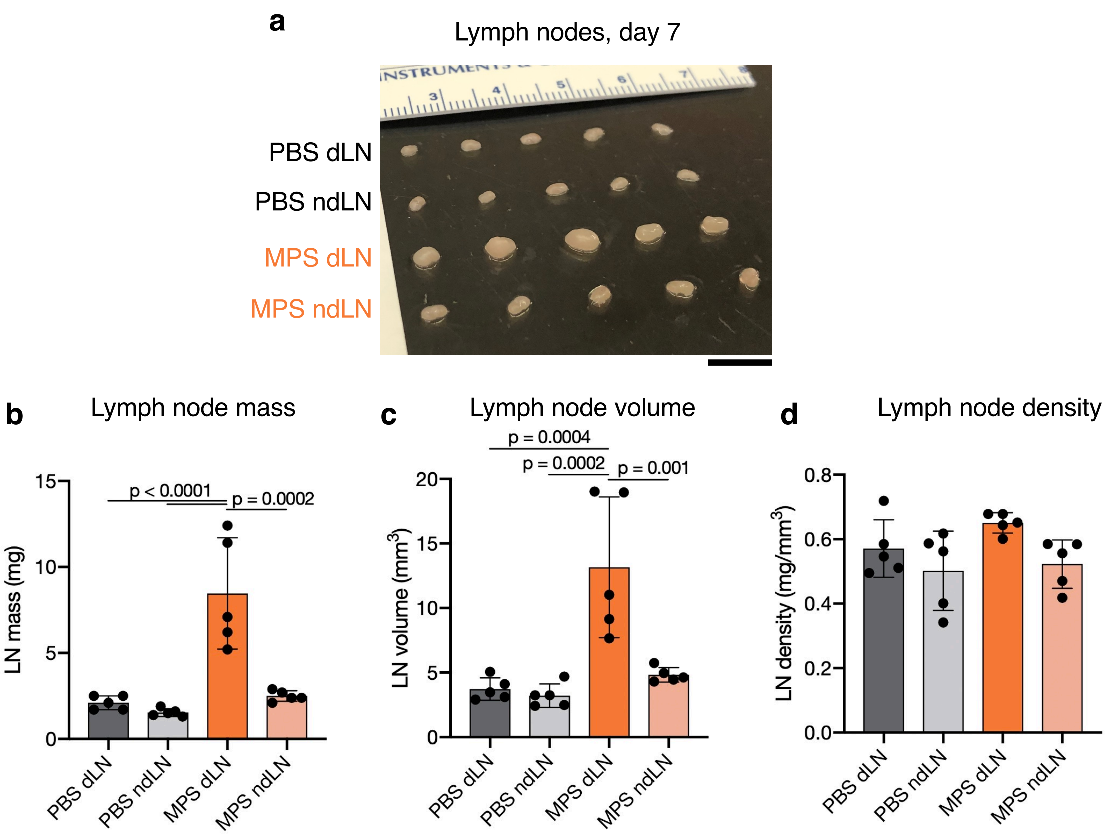


**Supplementary Figure 6. LN density after immunization.** Mice were immunized with MPS vaccines containing GM-CSF, CpG, and OVA protein and compared to control mice injected with PBS. Inguinal dLNs and ndLNs (contralateral to vaccine injection site) were harvested after 7 days. (a) Photograph of explanted LNs at day 7. LN mass (b), volume (c), and density (calculated as mass divided by volume) (d). For d, no statistically significant difference was found between groups. n = 5 biologically independent animals per group (MPS or PBS); means depicted; error bars, s.d. Statistical analysis was performed using analysis of variance (ANOVA) with Tukey's post hoc test. Scale bar = 1cm.

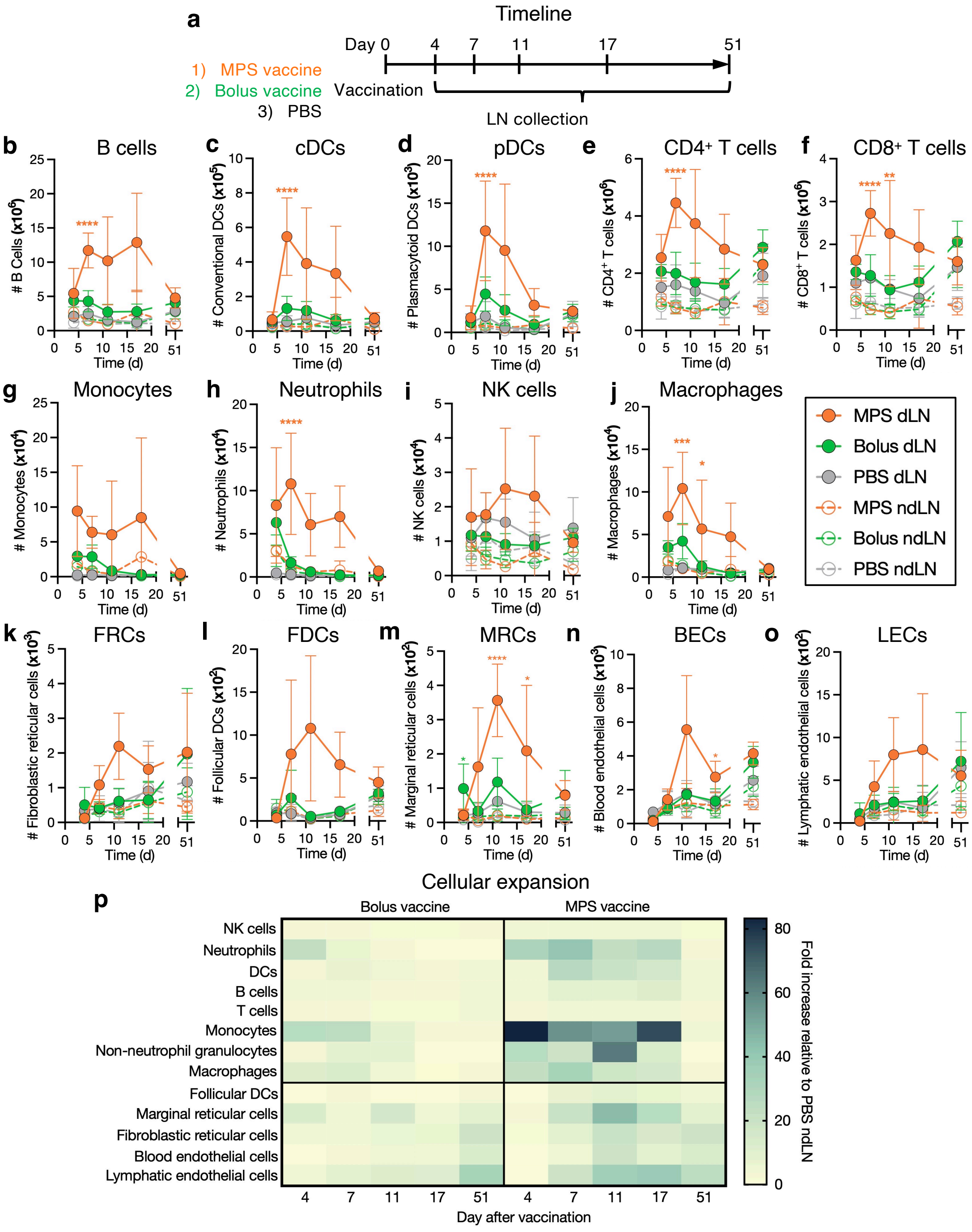


**Supplementary Figure 7. Immune and stromal cell populations in LNs after vaccination.** Mice were immunized with MPS or bolus vaccines containing GM-CSF, CpG, and OVA protein, euthanized on days 4, 7, 11, 17, and 51 for LN harvest and analysis through flow cytometry, and compared to PBS-injected controls. Immune and stromal populations were analyzed through flow cytometry over time. (a) Experimental timeline. (b) B cells (CD3$^-$ B220$^+$ CD11c$^-$ CD11b$^-$; re-plotted from Fig. 3c), (c) conventional DCs (CD3$^-$ B220$^-$ NK1.1$^-$ Ly6G$^-$ Ly6C$^-$ CD64$^-$ F4/80$^-$ CD11c$^+$ MHCII$^+$), (d)

plasmacytoid DCs (CD3$^-$ B220$^+$ CD11c$^+$ CD11b$^-$ Ly6C$^+$ Siglec-H$^+$), (e) CD4$^+$ T cells (CD3$^+$ B220$^-$ CD4$^+$ CD8$^-$), (f) CD8$^+$ T cells (CD3$^+$ B220$^-$ CD8$^+$ CD4$^-$), (g) monocytes (CD3$^-$ B220$^-$ NK1.1$^-$ Ly6G$^-$ CD11b$^+$ Ly6C$^+$), (h) neutrophils (CD3$^-$ B220$^-$ NK1.1$^-$ CD49b$^-$ CD11b$^+$ CD11c$^-$ SSC-A$^{int}$ Ly6G$^{hi}$), (i) natural killer cells (CD3$^-$ B220$^-$ NK1.1$^+$ CD49b$^+$), (j) macrophages (CD3$^-$ B220$^-$ NK1.1$^-$ Ly6G$^-$ Ly6C$^-$ CD64$^+$ F4/80$^+$), (k) fibroblastic reticular cells (CD45$^-$ CD31$^-$ PDPN$^+$ CD21/35$^-$ MAdCAM-1$^-$), (l) follicular dendritic cells (CD45$^-$ CD31$^-$ CD21/35$^+$; re-plotted from Fig. 3f), (m) marginal reticular cells (CD45$^-$ CD31$^-$ PDPN$^+$ CD21/35$^-$ MAdCAM-1$^+$), (n) blood endothelial cells (CD45$^-$ CD31$^+$ PDPN$^-$), and (o) lymphatic endothelial cells (CD45$^-$ CD31$^+$ PDPN$^+$) in LNs. (p) Heatmap of fold cellular expansion, relative to the PBS control, of the indicated cell types over time (days labeled below) after immunization with the bolus vaccine (left) or MPS vaccine (right). Mean cellular expansion was calculated relative to the PBS ndLN condition at day 4. For b-p, n = 4-5 biologically independent animals per group per timepoint; means depicted; error bars, s.d. Statistical analysis was performed using analysis of variance (ANOVA) with Tukey's post hoc test for normally distributed samples, and a Kruskal-Wallis test with Dunn's post hoc test otherwise; statistical significance is shown between the MPS dLN group and all other groups (*$P < 0.05$, ** $P < 0.01$, ***$P < 0.001$, ****$P < 0.001$).

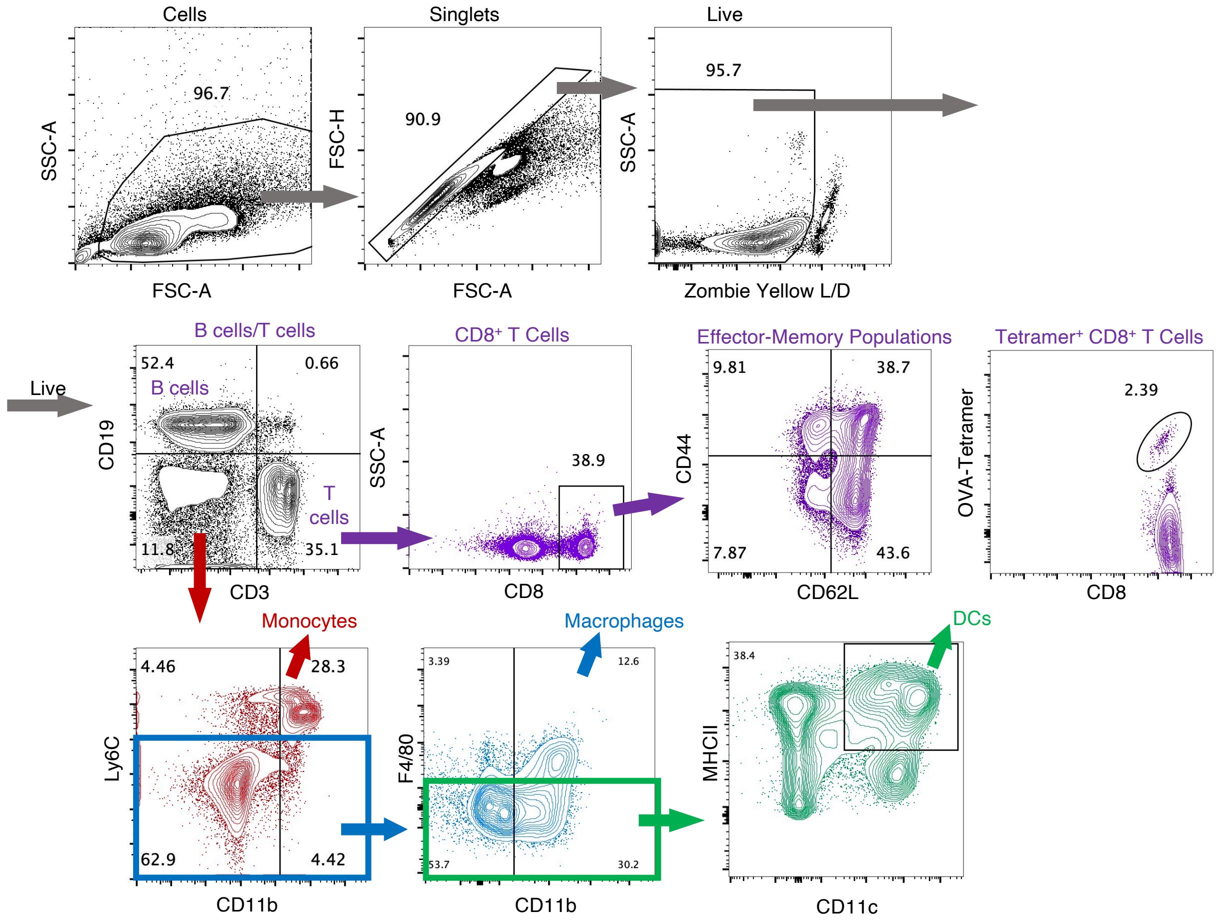


**Supplementary Figure 8.** Representative gating strategy to identify immune cell subsets.

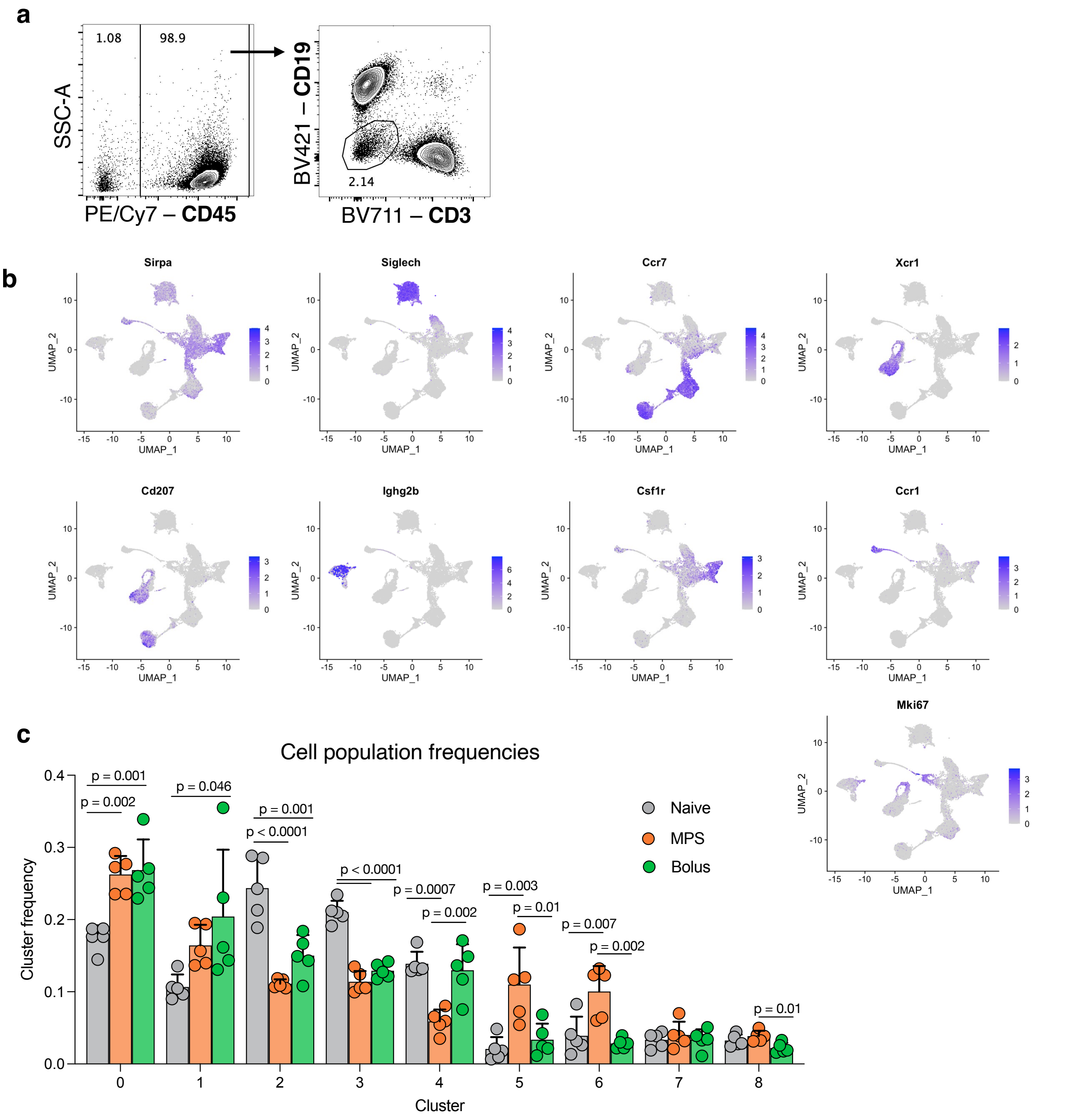


**Supplementary Figure 9. scRNAseq gating, cluster identification, and population frequencies.** (a) Representative flow cytometry plots depicting FACS-sorting strategy to identify myeloid cells ($CD45^{+}CD3^{-}CD19^{-}$) for scRNAseq analysis. (b) UMAP clustering of cells across conditions from Fig. 3h, with marker gene expression distinguishing the identified clusters (color scaled for each gene as indicated). (c) Frequency of individual cell clusters within each sample. Statistical analysis was performed using analysis of variance (ANOVA) with Tukey's post hoc test. n = 5 biologically independent animals per group.

**a**

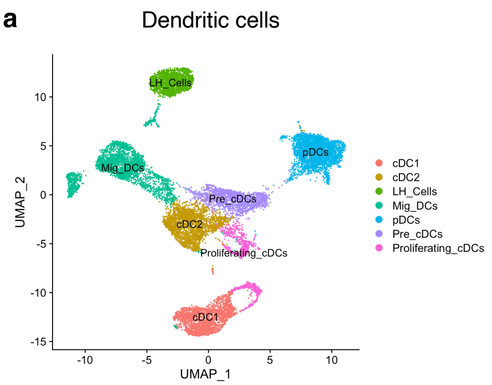


**b** **c**

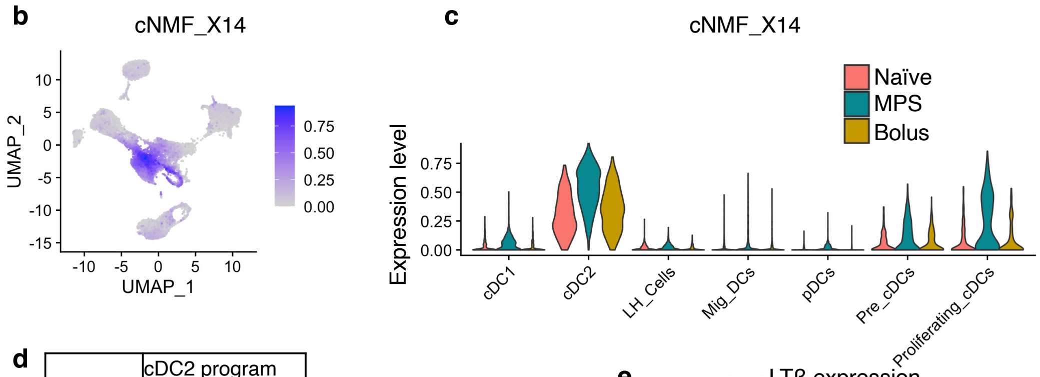


**d**

| | cDC2 program |
|---|---|
| Rank | cNMF_X14 |
| 1 | Ffar2 |
| 2 | Cass4 |
| 3 | Rasgef1b |
| 4 | Il1r2 |
| 5 | Elmo1 |
| 6 | Fam105a |
| 7 | Limd2 |
| 8 | Stap1 |
| 9 | Sirpb1a |
| 10 | Ltb |
| 11 | Ppp1r14a |
| 12 | Egr2 |

| | |
|---|---|
| 13 | Gm1966 |
| 14 | Gm6377 |
| 15 | Btg2 |
| 16 | Ifitm6 |
| 17 | Chn2 |
| 18 | Lst1 |
| 19 | Cd86 |
| 20 | Cyp4f16 |
| 21 | Bbs9 |
| 22 | Sirpa |
| 23 | Elmsan1 |
| 24 | Clec4a2 |
| 25 | Sirpb1b |

**e**

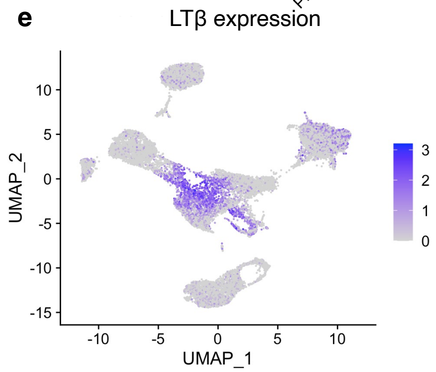


**Supplementary Figure 10. cDC2s are transcriptionally altered after MPS vaccination.** (a) UMAP re-clustering of dendritic cell populations from Fig. 3h. (b) Expression of cNMF X14 (cDC2-associated gene module) among DC populations, color scaled as indicated. (c) Expression level of cNMF X14 gene module among DC subsets. (d) Top 25 genes associated with cNMF X14 gene module. (e) Expression of *Ltb* among DC subsets.

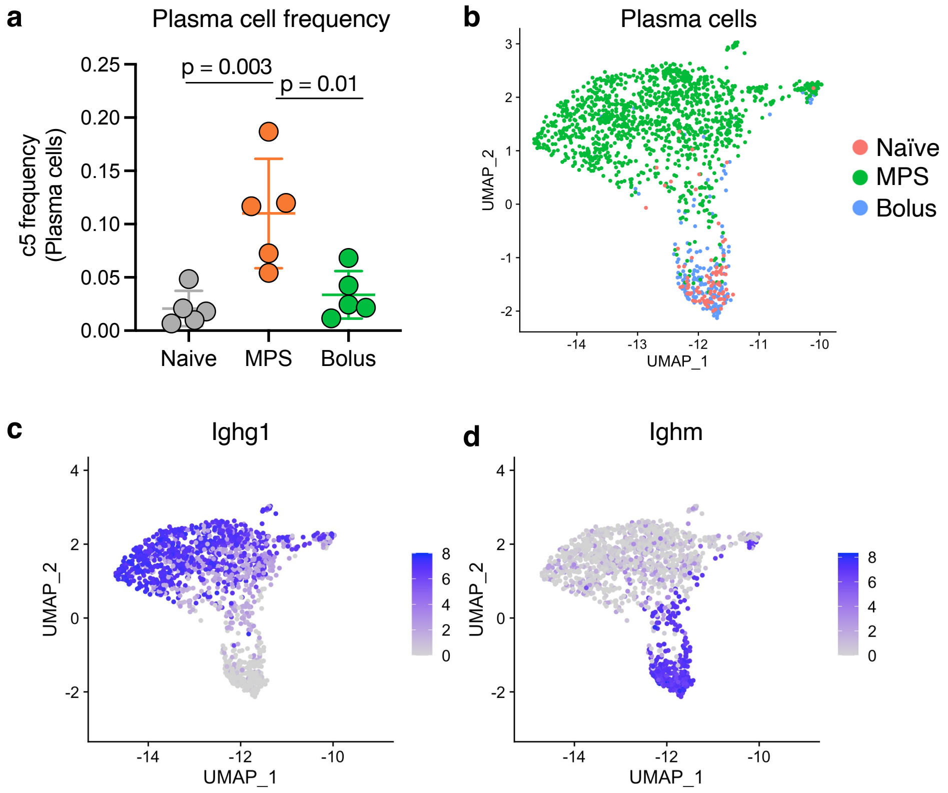


**Supplementary Figure 11. Plasma cells are enriched and express mature Ig after MPS vaccination.** (a) Frequency of plasma cells among different conditions. (b) UMAP re-clustering of plasma cells from Fig. 3h. Expression of (c) *Ighg1* and (d) *Ighm* among plasma cells, color scaled as indicated.

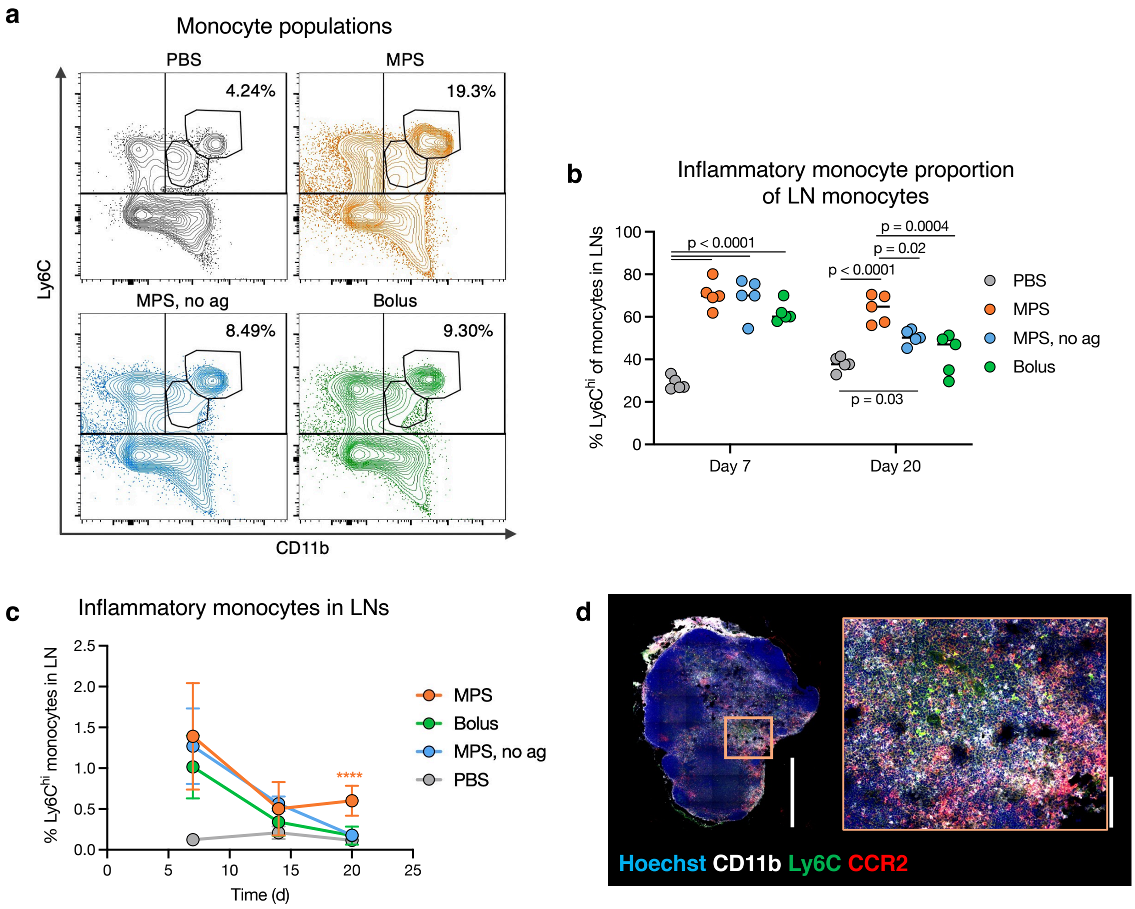


**Supplementary Figure 12. MPS vaccination expands inflammatory monocytes.** Mice were treated with MPS or bolus vaccines (containing GM-CSF, CpG, OVA), MPS vaccine without antigen (GM-CSF, CpG only), or PBS, and LNs were collected on days 7, 14, and 20 for cellular analysis. n = 5 biologically independent animals per group per timepoint. (a) Representative flow cytometry plots depicting $CD3^-$ $B220^-$ cells from LNs on day 20. Top right quadrant signifies monocytes ($CD11b^+$ $Ly6C^+$); upper population in that quadrant signifies inflammatory $Ly6C^{hi}$ monocytes. (b) Proportion $Ly6C^{hi}$ of monocytes ($CD3^-CD19^-CD11b^+Ly6C^+$) in LNs on days 7 and 20. (c) $Ly6C^{hi}$ inflammatory monocyte proportions in the LN over time. (d) IHC image depicting an MPS-vaccinated mouse LN extracted on day 20 and stained for inflammatory monocyte markers. Scale bars = 0.5mm (left), 100µm (right). For b-c, statistical analyses were performed using analysis of variance (ANOVA) with Tukey's post hoc test; means depicted; error bars, s.d. For c, only differences present between one group and all other groups are shown (*P < 0.05, ** P < 0.01, ***P < 0.001, ****P < 0.001).

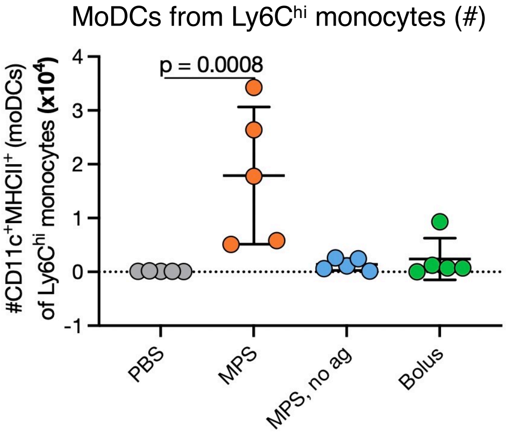


**Supplementary Figure 13. Monocyte-derived DCs in LNs.** Mice were treated with MPS or bolus vaccines (containing GM-CSF, CpG, OVA), MPS vaccine without antigen (GM-CSF, CpG only), or PBS, and LNs were collected on day 20 for cellular analysis. Number of monocyte-derived DCs from $Ly6^{hi}$ inflammatory monocytes ($CD3^{-}B220^{-}CD11b^{+}Ly6C^{hi}CD11c^{+}MHCII^{+}$ cells) in the LN at day 20. Statistical analysis was performed using a Kruskal-Wallis test with Dunn's post hoc test. Means depicted; error bars, s.d.; n = 5 biologically independent animals per group.

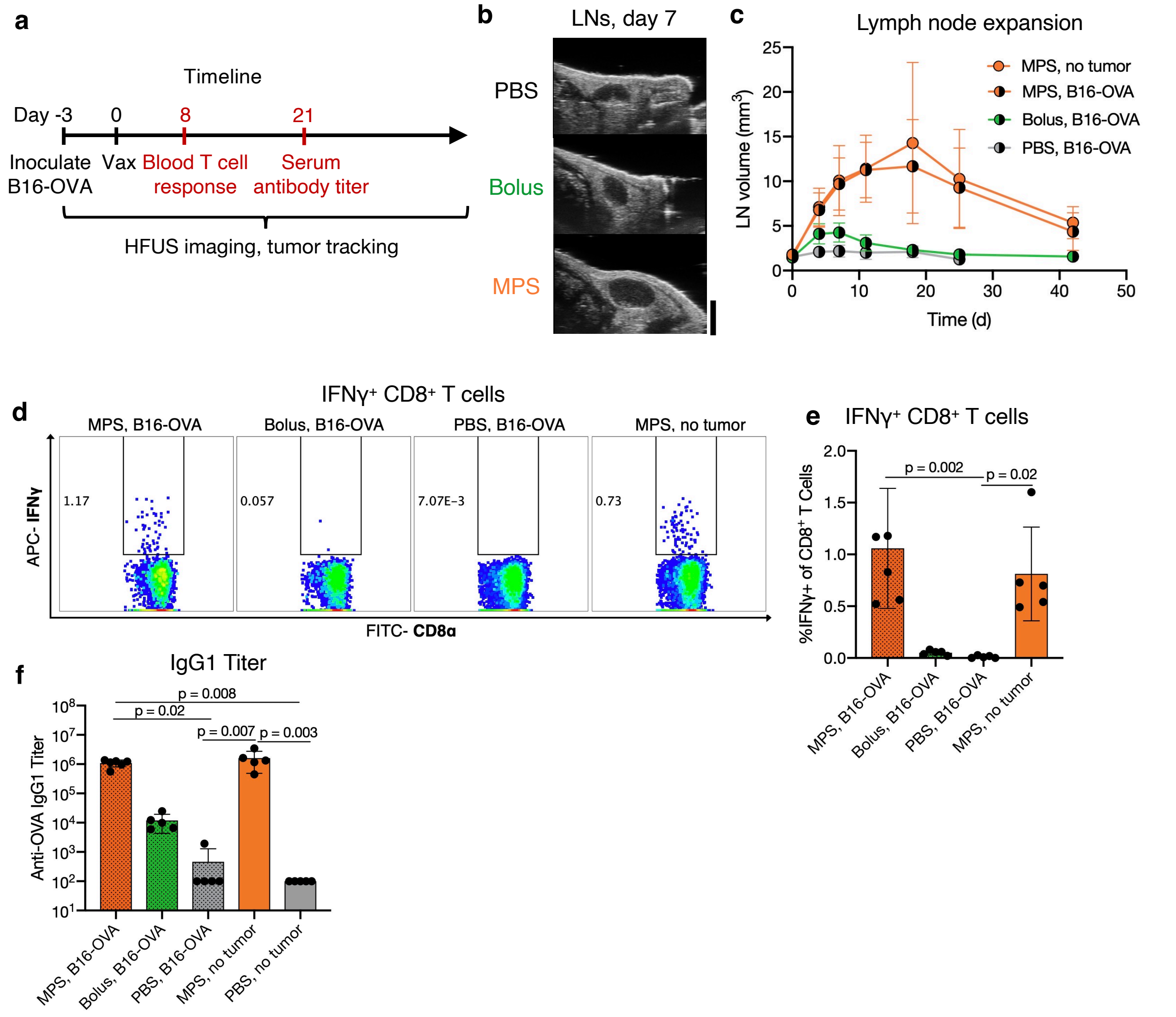


**Supplementary Figure 14. Therapeutic study to assess correlations of LN expansion with vaccine efficacy.** Mice were inoculated with B16-OVA tumors and three days later treated with MPS or bolus vaccines containing GM-CSF, CpG, and OVA protein, and compared to PBS-injected controls. A fourth group of tumor-free mice was treated with MPS vaccines (called “MPS, no tumor”). Inguinal dLNs were imaged using HFUS at multiple timepoints, and blood was collected 8 and 21 days after vaccination to assess T cell responses and serum antibody titers, respectively. n = 5-6 biologically independent animals per group. (a) Experimental timeline. (b) Representative images of dLNs 7 days after vaccination. Scale bar = 2mm. (c) Quantification of LN volume over time. No PBS-injected mice survived at day 42. (d) Representative flow cytometry plots of IFNγ$^+$ CD8$^+$ T cells after SIINFEKL peptide restimulation. (e) Proportion IFNγ$^+$ of CD8$^+$ T cells in blood after SIINFEKL peptide restimulation. Statistical analysis was performed using a Kruskal-Wallis test with Dunn’s post hoc test. (f) Serum anti-OVA IgG1 antibody titer, 21 days after immunization. Statistical analysis was performed using a Kruskal-Wallis test with Dunn’s post hoc test. For c and d, means are depicted; error bars, s.d.

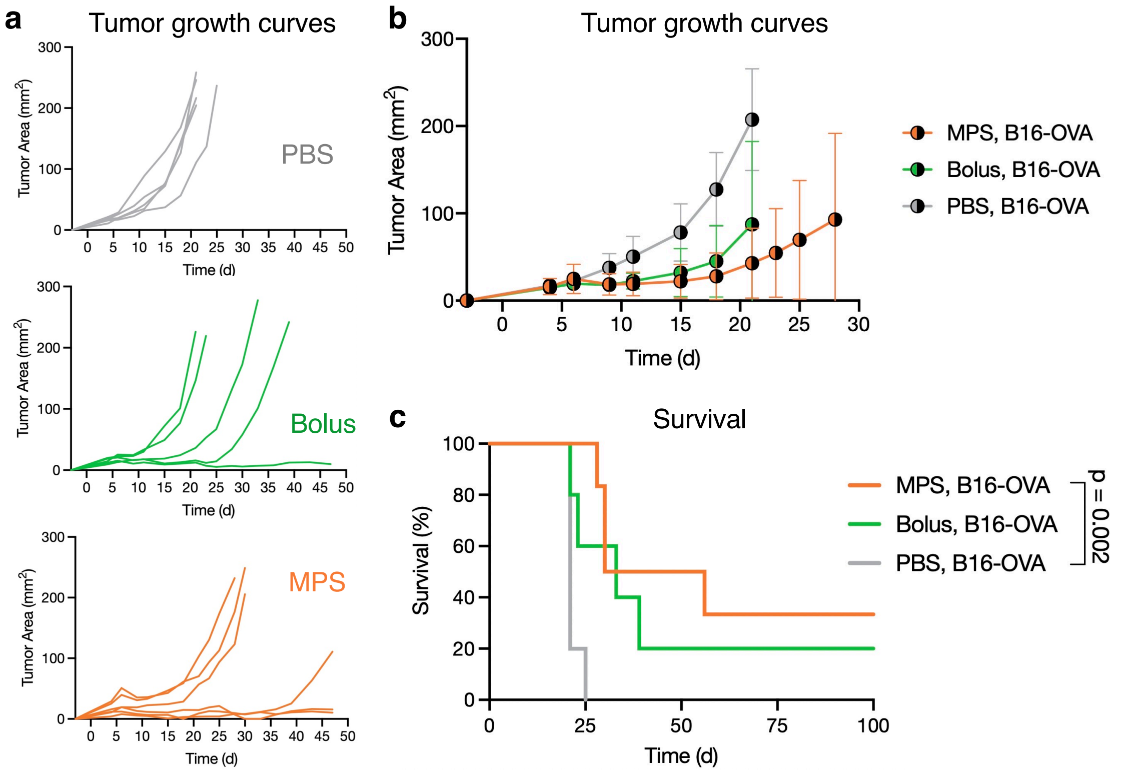


**Supplementary Figure 15. Vaccine therapeutic efficacy.** Mice were treated as described in Supplementary Fig. 4. Tumor growth was tracked externally using calipers and mice were euthanized at humane endpoints. (a) Individual growth curves of treated mice. (b) Combined growth curves. (c) Kaplan-Meier curves depicting survival of groups. n = 5-6 biologically independent animals per group; statistical analysis was performed using a log-rank (Mantel-Cox) test, correcting for multiple comparisons.

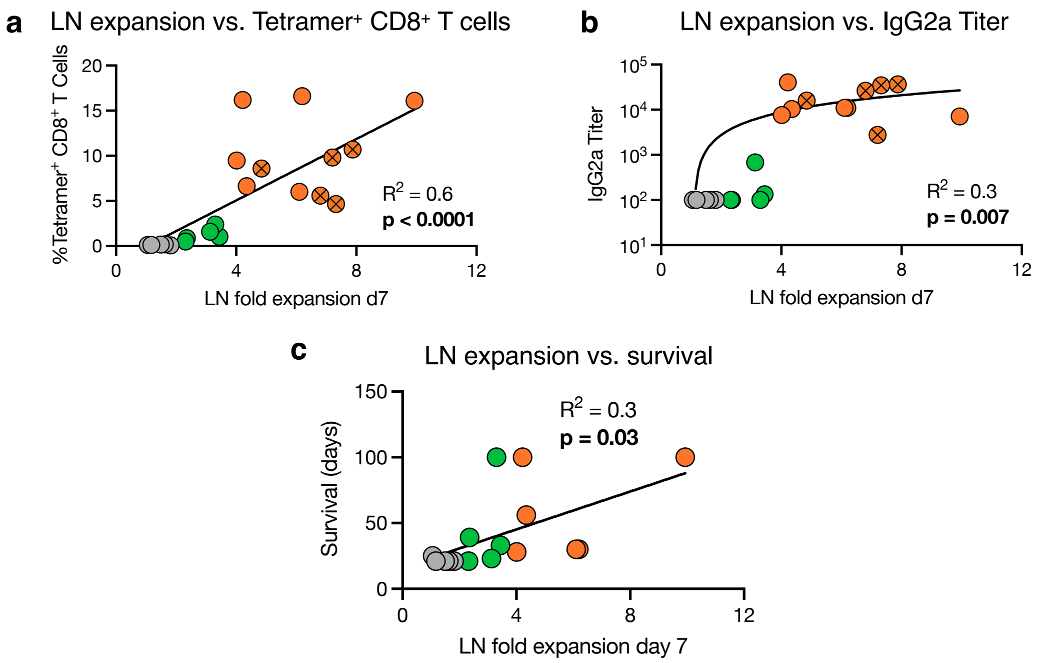


**Supplementary Figure 16. The adaptive, antitumor vaccine response correlates with LN expansion.** LN expansion from Fig. 6b is plotted against vaccine response data from Fig. 6 and Supplementary Fig. 18. Linear regressions are shown of LN fold expansion 7 days after vaccination versus (a) OVA-tetramer+ CD8+ T cells, (b) anti-OVA IgG2a titers, and (c) long-term survival. For b, the y-axis is plotted on a log scale to show the range of vaccine responses, although a linear regression was performed. Correlations are statistically significant.

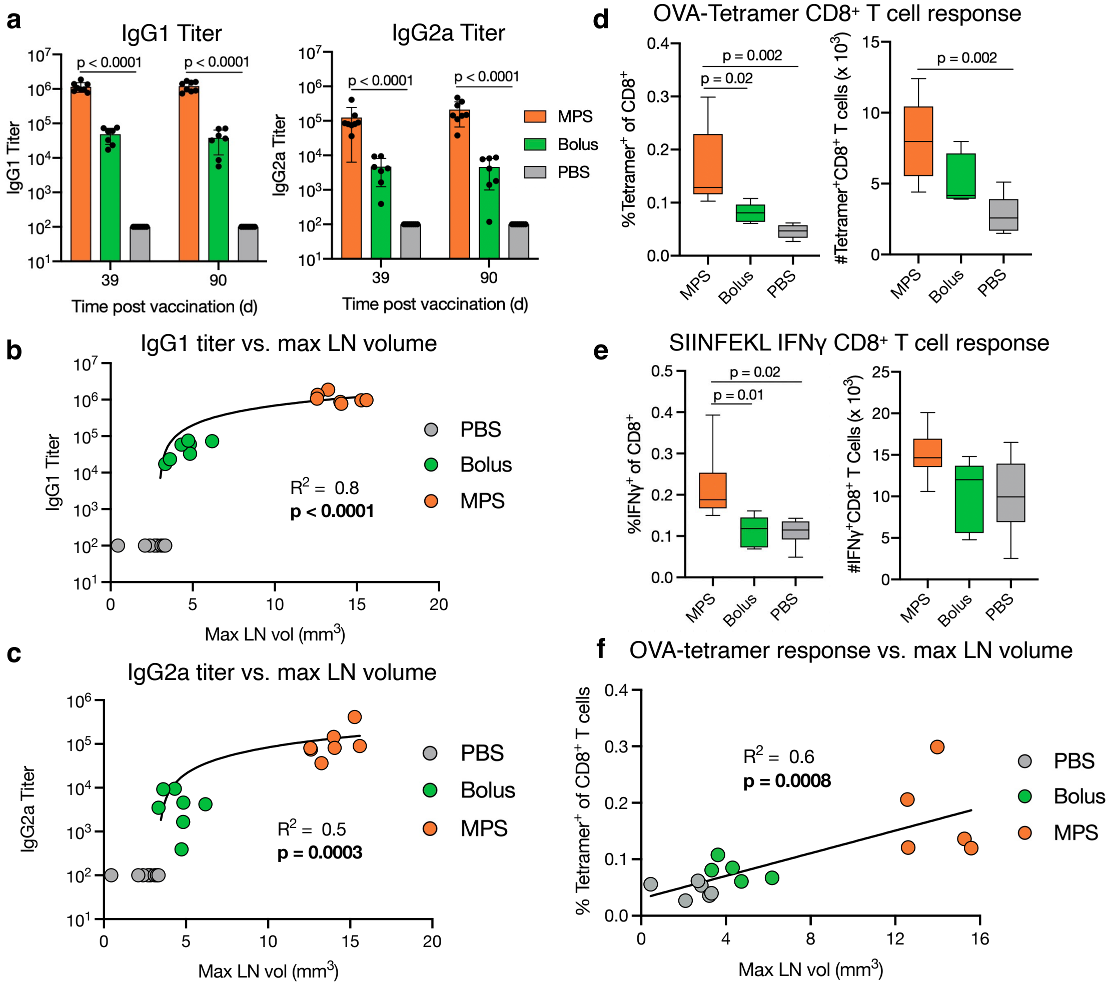


**Supplementary Figure 17. Long-lived vaccine responses correlate with LN expansion.** Mice were immunized with MPS or bolus vaccines delivering GM-CSF, CpG, and OVA protein (same setup as Fig. 1). 39 and 90 days after vaccination, blood was collected for OVA-specific serum antibody titer analysis using ELISA. 103 days after vaccination, mice were euthanized and spleens were collected for T cell analysis. (a) Serum IgG1 (left) and IgG2a (right) titers against OVA. n = 7-8 biologically independent animals per group; means depicted; error bars, s.d. Statistical analysis was performed using a Kruskal-Wallis test with Dunn's post hoc test. Linear regressions (plotted on log y-axis) of IgG1 titer (b) and IgG2 titer (c) versus the maximum LN volume in each mouse. Correlations are statistically significant. Proportion (left) and number (right) of OVA-tetramer⁺ CD8⁺ T cells (d) and IFNγ⁺ CD8⁺ T cells (e) in spleens. For d-e, n = 5-6 biologically independent animals per group. Whiskers extend min-to-max with box displaying 25$^{th}$ to 75$^{th}$ percentiles. Statistical analysis was performed using analysis of variance (ANOVA) with Tukey's post hoc test (d, e right) or Kruskal-Wallis test with Dunn's post hoc test (e left). (f) Linear regression of OVA-tetramer$^+$ CD8$^+$ T cell proportion with maximum LN volume.

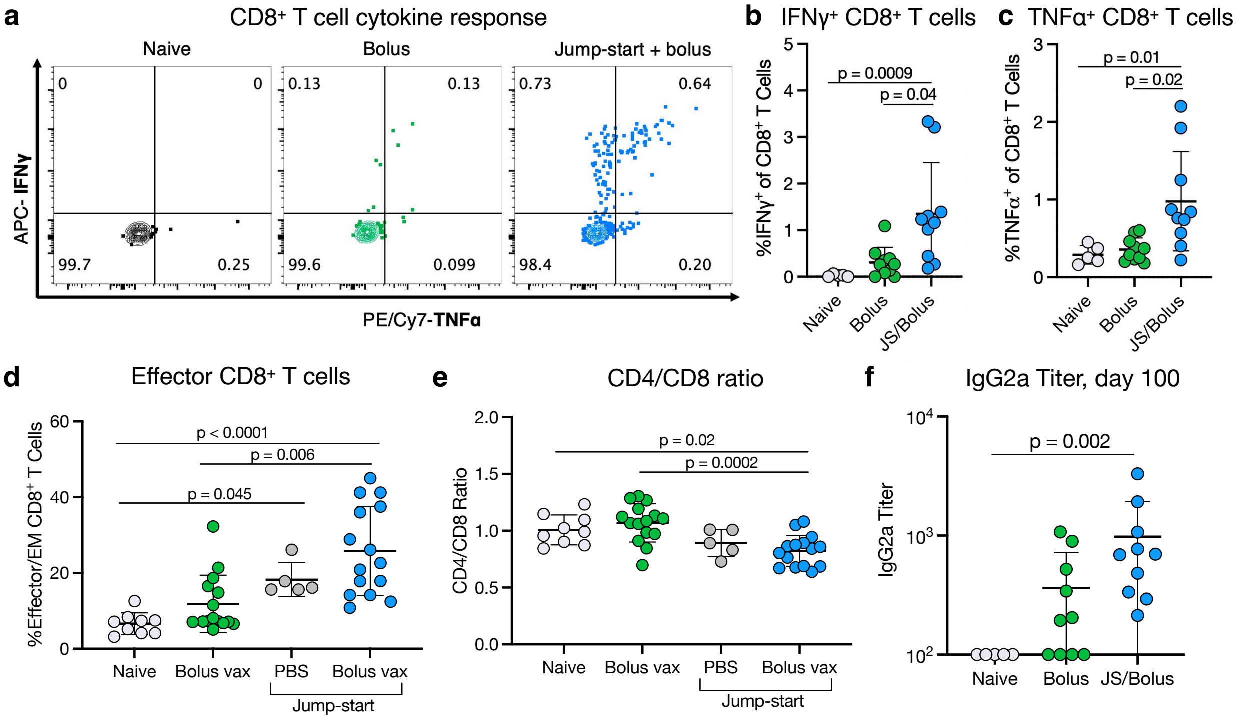


**Supplementary Figure 18. Antigen-free MPS "jump-start" strategy boosts bolus vaccine response.** Mice were injected with PBS or a bolus vaccine on day 0, or injected with an MPS no-antigen "jump-start" on day -7 followed by PBS or a bolus vaccine (GM-CSF, CpG, and OVA protein) on day 0. Mice were bled after 8 and 100 days for T cell analysis and serum antibody titers, respectively. (a) Representative flow cytometry plots depicting $CD8^+$ T cell cytokine production after restimulation with OVA SIINFEKL peptide. Quantification of IFNγ (b) and TNFα production (c). (d) Proportion of effector-phenotype ($CD44^+CD62L^-$) $CD8^+$ T cells in the blood. Statistical analysis was performed using a Kruskal-Wallis test with Dunn's post hoc test. (e) CD4/CD8 T cell ratio in the blood. Statistical analysis was performed using analysis of variance (ANOVA) with Tukey's post hoc test. (f) Anti-OVA IgG2a titer at 100 days. For b-c and f, n = 5-10 biologically independent animals per group; statistical analysis was performed using a Kruskal-Wallis test with Dunn's post hoc test. For d-e, n = 10-15 biologically independent animals per group; results are combined from two independent experiments. For b-e, means depicted; error bars, s.d.

**a**

Experimental groups

| Group | Day -7 | Day 0 |
|---|---|---|
| Naïve | n/a | n/a |
| Bolus | n/a | Bolus vaccine |
| Jump-start/naïve | MPS, no ag | n/a |
| Jump-start/bolus | MPS, no ag | Bolus vaccine |

**b**

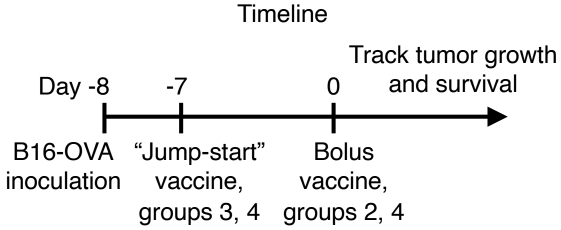


**Supplementary Figure 19. Therapeutic "jump-start" experiment layout.** Mice bearing B16-OVA tumors (day -8) were treated at day -7 with an MPS "jump-start" (MPS material, GM-CSF, and CpG without antigen) or left untreated and were injected with a bolus vaccine (GM-CSF, CpG, and OVA) or left untreated. Tumor growth and survival were tracked. (a) Experimental groups. (b) Timeline.
